# Retracing the evolution of a modern periplasmic binding protein

**DOI:** 10.1101/2023.05.30.542879

**Authors:** Florian Michel, Sergio Romero-Romero, Birte Höcker

## Abstract

Investigating the evolution of structural features in modern multidomain proteins helps to understand their immense diversity and functional versatility. The class of periplasmic binding proteins (PBPs) offers an opportunity to interrogate one of the main processes driving diversification: the duplication and fusion of protein sequences to generate new architectures. The symmetry of their two-lobed topology, their mechanism of binding, and the organization of their operon structure led to the hypothesis that PBPs arose through a duplication and fusion event of a single common ancestor. To investigate this claim, we set out to reverse the evolutionary process and recreate the structural equivalent of a single-lobed progenitor using ribose-binding protein (RBP) as our model. We found that this modern PBP can be deconstructed into its lobes, producing two proteins that represent possible progenitor halves. The isolated halves of RBP are well folded and monomeric proteins, albeit with a lower thermostability, and do not retain the original binding function. However, the two entities readily form a heterodimer *in vitro* and *in-cell*. The X-ray structure of the heterodimer closely resembles the parental protein. Moreover, the binding function is fully regained upon formation of the heterodimer with a ligand affinity similar to that observed in the modern RBP. This highlights how a duplication event could have given rise to a stable and functional PBP-like fold and provides insights into how more complex functional structures can evolve from simpler molecular components.

## Introduction

The detection of chemicals in the environment, their molecular recognition and transport into the cell as well as the resulting downstream signaling is an integral part of life in any cell. As one of the central classes of proteins responsible for this function in prokaryotes, the periplasmic binding proteins (PBPs) serve as an important element in these complex response networks (Matilla, 2021). These bilobal proteins are involved in the transport of a wide variety of substrates, and are generally considered to belong to an ancient protein fold (Clifton, 2016; Felder, 1999).

The PBP architecture consists of two opposing lobes, with each lobe being built of a central, five-stranded parallel β-sheet with five α-helices flanking its sides. The two lobes are connected via a hinge region, with the complexity and number of crossovers dependent on the class of PBP. This architecture also gives rise to the most common mechanism in which PBPs recognize and bind their ligands (Chandravanashi, 2021; Scheepers, 2016; Berntsson, 2010). This distinct mode of binding that a majority of PBPs follow is a “venus flytrap-like’’ mechanism and considered one of the hallmark features of this protein class (Felder, 1999). While in the unbound state, PBPs are in an “open” form with a space created by the two lobes accessible to surrounding solutes. Recognition and binding of the ligand facilitates interaction between the two lobes, leading to the eponymous hinge-bending motion which results in the “closed” conformation with the cleft now being tightly shut around the ligand, excluding the solvent upon binding (Berntsson, 2010; Felder, 1999). This common binding mechanism is reflected in PBPs that bind similar molecules with very different selectivities and affinities at the same binding site (Kröger, 2021a). For these reasons, PBPs have been used in several engineering and design approaches, especially creating highly sensitive biosensors and molecular switches (Steffen, 2016; Jeffery, 2011; Medintz, 2006; Dwyer, 2004), and designing new binding properties (Banda-Vázquez, 2018; Scheib, 2014, Kröger, 2021b).

Despite diversity in the sequences of different PBPs, a shared common ancestry has been proposed a while ago (Fukami-Kobayashi, 1999; Louie, 1993). Their structural features, similarities in binding mechanism, and shared operon structure – with the PBP being on the same operon as the associated signaling proteins downstream – have long led to the theory that PBPs arose via gene duplication of a progenitor protein and subsequent diversification. However, it is unclear in which order these events might have occurred (Fukami-Kobayashi, 1999). It has been previously suggested that this common ancestor could have been a CheY-like protein adopting a flavodoxin-like fold. Formation of an ancestral dimer in combination with a gene duplication and fusion event might have led to the typical bilobal structure of the modern PBP (Fig. 1A), an event that has already been investigated for the evolution of other protein folds (Toledo-Patiño, 2019; Farías-Rico, 2014).

**Figure 1.**
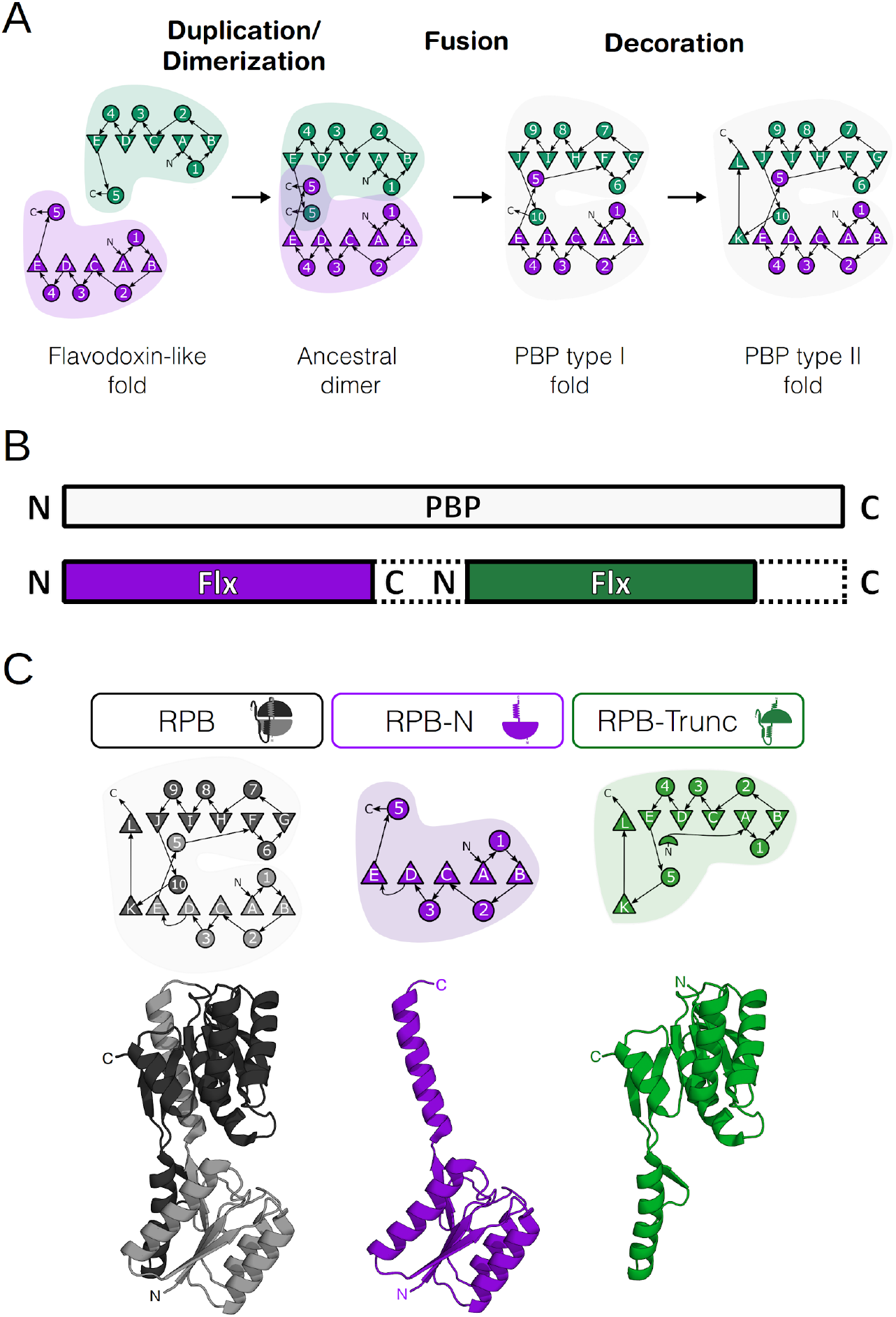
Proposed evolutionary trajectory of modern PBPs and the derived constructs used in this study. **(A)** Proposed steps that reconstruct the evolution of modern periplasmic-binding-protein (PBP) folds from an ancestral protein adapting the flavodoxin-like fold (adapted from Fukami-Kobayashi, 1999). A duplication and dimerization along with swaps in secondary structure led to the formation of an ancestral dimer. Subsequent fusion of the genes then led to the emergence of the Type I PBP-like fold and further changes of secondary structure to that of the Type II PBP-like fold. **(B)** Schematic representation of the profile-profile alignments for a representative full-length PBP with the flavodoxin-like-fold (Flx). **(C)** First-generation constructs RBP (black), RBP-N (violet) and RBP-Trunc (green) analyzed in this work to recreate the PBP-halves.

Although the sequences of modern PBPs have diversified from their evolutionary ancestors, the topology is predominantly conserved. There are mainly two classes of PBP, with a slight difference in the order of secondary structural elements. It is thought that the second class descends from already evolved class I PBPs even though sequence similarity is not high between the two folds (Fukami-Kobayashi, 1999). They are in fact classified as independent folds of either PBP-like I or PBP-like II in SCOP (Chandonia, 2019), as being of the same topology level as flavodoxins (for type I) and an independent homology group (for type II) in ECOD (Cheng, 2015), and as two different superfamilies in CATH (Sillitoe, 2021). The application of modern bioinformatic resources has opened up new opportunities to revisit some of these concepts of evolutionary relationships, partially through emergence of tools to more efficiently probe sequence space also in the sub-domain regime of proteins (Farías-Rico, 2014; Ferruz, 2020; Nepomnyachiy, 2017; Alva, 2015).

In this work we combine the approach of a sequence profile-profile comparison analysis using Hidden-Markov-Models (HMMs) with a structural comparison of the two lobes of the PBP-like fold type I. Based on this analysis, the emergence of the PBP-like fold via the duplication of a flavodoxin-like ancestor can be revisited. To further substantiate the claim, we biophysically and structurally characterized truncated constructs of the ribose-binding protein from *Thermotoga maritima* (RBP) that correspond to the proposed duplicated progenitor halves. We found that it is generally possible to obtain stable and well folded monomeric proteins expressing only the individual lobes of full-length RBP. The two independent halves appear to readily form a heterodimer, while also reconstituting the ribose-binding ability of the parental protein, with affinities in the same order of magnitude. These results suggest a plausible path for the evolution of modern PBPs and increase our understanding of the evolution of complex and multidomain proteins from smaller molecular components.

## Results and Discussion

### Disassembling a modern RBP into likely progenitor halves

The proposed mechanism of a duplication event being responsible for the architecture of PBPs mostly relies on analysis of either the available structures of modern PBPs (Poolman, 2010; Louie, 1993), or comparison of the sequences of PBP-like and flavodoxin-like proteins (Fukami-Kobayashi, 1999). We wanted to investigate whether the duplication of the flavodoxin-like progenitor is not only theoretically feasible, but also practically. To retrace the evolution of a PBP, we characterized constructs based on the halves of a ribose-binding protein (Fig. 1 and Table S1). This not only allows to probe the plausibility of this mechanism in general, but also offers an opportunity to investigate the individual impact of each subdomain-part on the stability and function of modern PBPs.

We chose the Ribose-binding-protein (RBP) of *T. maritima* for this purpose. Not only does the thermophilic nature of this protein offer a robust system, but also a previously reported expression of a 21 kDa truncated version (Cuneo, 2008) made this an excellent candidate for a model system. To generate an overview of possible intersections, a multiple sequence alignment with RBP as input was generated with HHpred (Fig. S1). The results show not only the alignment of other full-length PBPs on the query sequence but also an alignment of the individual lobes. The lobes align with a clear cut being observable between residue 30-155 and 156-310 of the RBP (numbering consistent with uniprot entry Q9X053). To compare this with the alignment of the proposed progenitor flavodoxin-like proteins, the same alignment was generated within the *Fuzzle* database (Ferruz, 2021), which automatically excludes sequences of the same fold. It shows that flavodoxin-like proteins align with both the corresponding N- and C-terminal halves of the PBP sequence (Ferruz, 2021). While alignment of flavodoxin-like proteins with RBP seems to heavily favor hits on the N-terminal half, some hits are also found with the C-terminal half. A reason why less hits might be observed on the C-terminal half of this modern RBP could be a result of the duplication and a subsequent decoupling of the sequences of the two halves, resulting in increased divergence from the progenitor flavodoxin-like protein, and thereby making it harder to identify.

While the existence of the earlier reported truncated RBP variant could be an artifact of the expression in *Escherichia coli* (Cuneo, 2008), it is also possible to be a natural occurrence. A shortened version of a solute binding protein with a proposed biological function has been reported previously (Bae, 2018). Although it is unclear why these single-lobed proteins might exist, the truncated RBP could also carry biological significance. Thus, we chose to use the truncated protein that is roughly the equivalent of the single-lobed half as a base for the constructs used in this study.

For the first generation of constructs we took to the lab, the sequence of the full-length RBP was disassembled into the corresponding halves (Fig. 1 and Table S1). The site of dissection was determined by structural alignment of RBP in absence of ribose (PDB ID: 2FN9) to the top scoring flavodoxin-like proteins in the HHpred analysis, resulting in the constructs RBP-N (amino acid 30-153 of RBP) and RBP-C (amino acid 157-291). These constructs were expressed and characterized using biochemical and biophysical methods.

### RBP halves are well folded

Upon overexpression of the RBP halves in *E. coli* the protein RBP-N was found in the soluble fraction of the cell extract while RBP-C was located in inclusion bodies. Since full-length RBP also features a C-terminal decoration common to modern PBPs which does not correspond to any elements in the canonical flavodoxin-like architecture, the additional elements (two β-strands that facilitate another cross-over between the two lobes and extend the central β-sheet of the two halves) had been removed in RBP-C. This removal might be the reason why in contrast to RBP-N, which expressed solubly, could be purified to homogeneity, and remained stable at concentrations above 15 mg mL^-1^, RBP-C only expressed insolubly. We therefore decided to continue the investigation with the truncated construct RBP-Trunc (residues 142-310) instead (Fig. 1C), which is related to the RBP-C half and expressed solubly with similar stability to the N-terminal construct RBP-N.

Both RBP-N and RBP-Trunc display far-UV CD spectra with the signature ɑ-helix minima at 208 and 222 nm and moderated by the signal of the β-sheet at 218 nm, both characteristic for α/β-proteins (Fig. 2A) and comparable with the native full-length RBP. Comparison of the intrinsic fluorescence also corroborates this (Fig. S2A), indicating that the constructs are well folded since the intensity maximum suggests that the aromatic residues are buried from the solvent.

**Figure 2.**
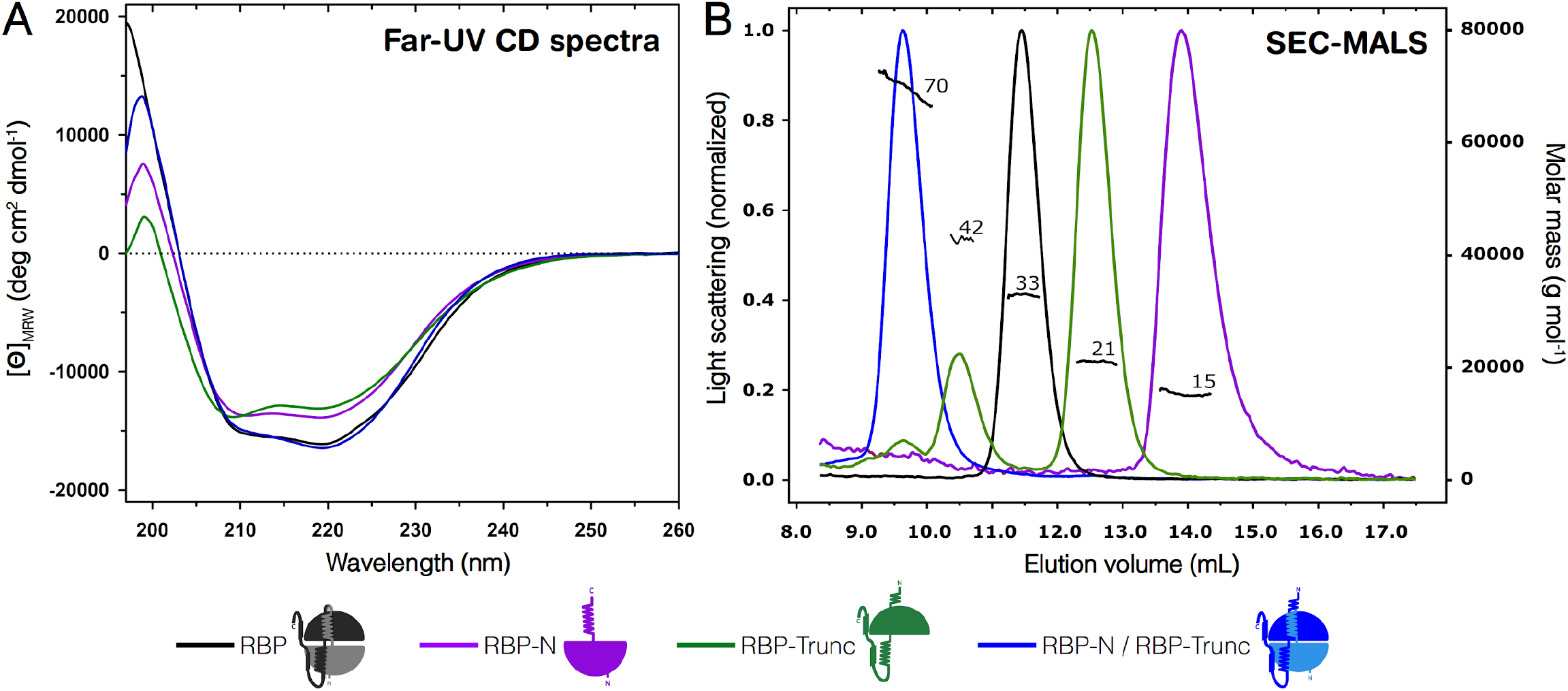
Biophysical characterization of the first-generation constructs. **(A)** Far-UV CD spectra at 20 °C collected in 10 mM sodium phosphate, 50 mM sodium chloride, pH 7.8. **(B)** SEC-MALS experiments performed in 10 mM sodium phosphate, 50 mM sodium chloride, 0.02% sodium azide, pH 7.8. Numbers indicate the molecular weight determined after data analysis. Values derived from the experiments are reported in Supplementary Table S2. In both panels, the color code is RBP (black), RBP-N (violet), RBP-Trunc (green) and the RBP-N/RBP-Trunc dimer (blue).

Further analysis with SEC-MALS (Fig. 2B) confirmed the monomeric state of RPB and RBP-N. However, RBP-Trunc is in an equilibrium of mostly monomeric species and dimer, with higher oligomers also being present (Table S2). These results indicate that the RBP halves are well folded proteins and express mainly as monomeric systems, similar to those observed in another PBP, HisJ (Chu, 2013). To follow up on this, we continued to study their properties in the presence of each other.

### RBP halves form a heterodimer whose structure is identical to full-length RBP

Since one of the steps proposed in the evolution of the modern PBP architecture involves an ancient dimer, we investigated whether the obtained constructs had the ability to reconstitute the full-length RBP fold. For this, the individually purified RBP-N and RBP-Trunc were mixed in an equimolar ratio and then analyzed. The far-UV CD spectra (Fig. 2A) show a significant change of the signal to the individual constructs, with the signal of the mixed RBP-N/RBP-Trunc resembling that of the full-length RBP. A similar behavior can be observed in the intrinsic fluorescence spectra (Fig. S2A), where the original characteristics of the full-length protein are reconstituted when mixed *in vitro*, hinting at the formation of an RBP-N/RBP-Trunc heterodimer. Complex formation is supported by SEC-MALS analysis where only one well-defined peak is displayed corresponding to the mass of the RBN-N/RBP-Trunc dimer (Fig. 2B and Table S2).

Additionally, DSC analysis of the proteins supports the formation of a dimer that resembles the parental protein. All endotherms show clear single and cooperative transitions, as has been observed for other PBPs such as maltose-, arabinose-, and histidine-binding proteins (Kreimer, 2003; Ganesh, 1997; Fukada, 1983). However, RBP and its halves showed irreversible thermal unfolding possibly due to their thermophilic nature, contrary to most PBPs which exhibit reversible transitions (Vergara, 2020; Aggarwal, 2011; Prajapati, 2007; Kreimer, 2003; Ganesh, 1997; Fukada, 1983). While full-length RBP has a *T_m_* of 106.9 °C similar to the one previously reported for the construct (Cuneo, 2008), RBP-N and RBP-Trunc show lower thermostability with a *T_m_* of 76.6 °C and 73.3 °C, respectively (Fig. 3A, Table 1, and Table S3). The results show that the halves have native-like properties, that interdomain interactions are important in RBP and that these provide relevant stabilization, in the same way as has been described for other multidomain proteins (Vergara, 2023; Liu, 2019; Kantaev, 2018; Vogel, 2004; Wenk, 1998; Careaga, 1995; Brandts, 1989). This decrease in thermostability of the individual constructs is compensated by the formation of the RBP-N/RBP-Trunc dimer, whose *T_m_* is shifted by more than 20°C to 99.7 °C, more closely resembling that of RBP.

**Figure 3.**
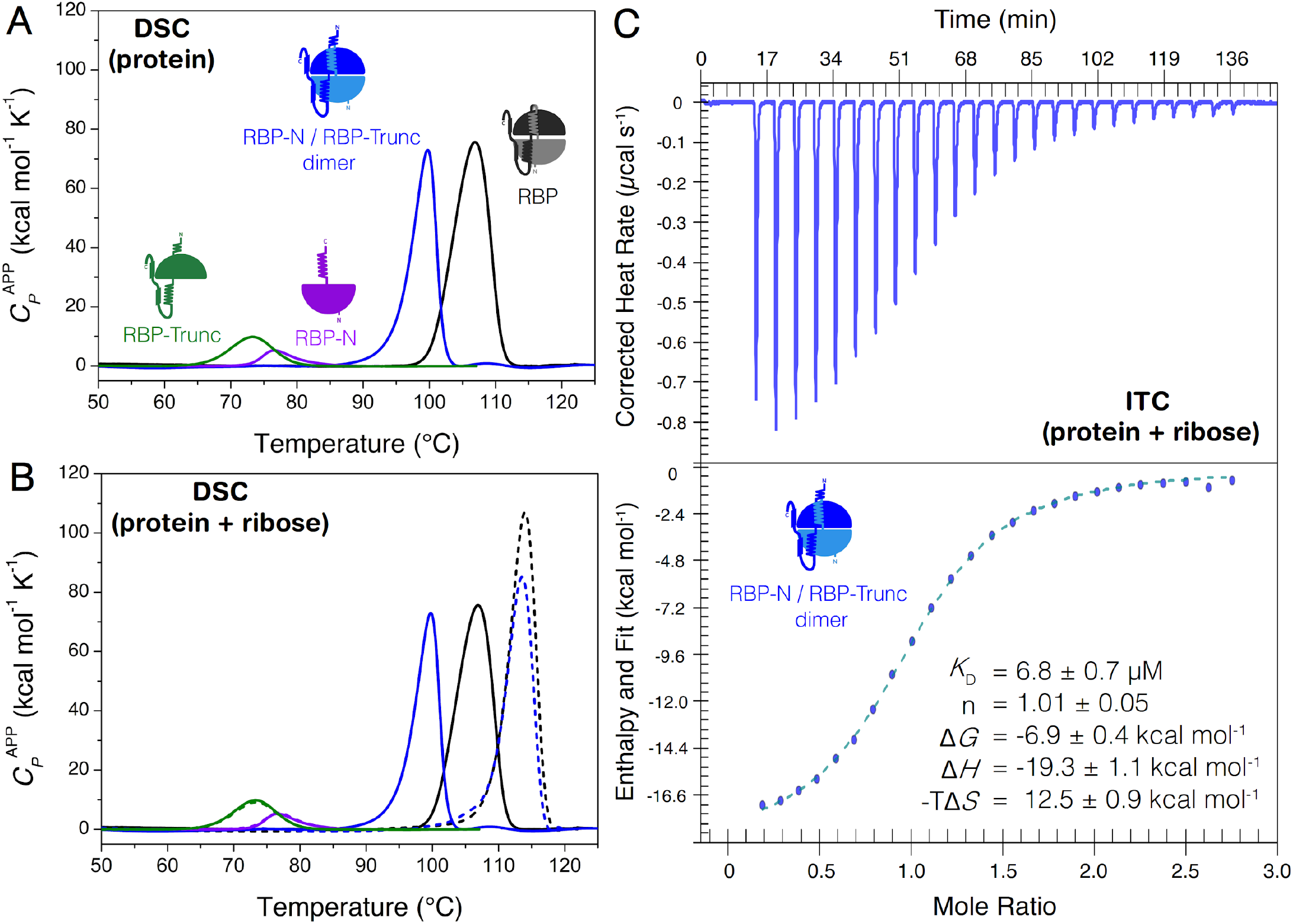
Thermodynamic characterization of the first-generation constructs and their interaction with ribose. **(A)** DSC endotherms at 1.5 °C min^-1^ of the halves RBP-Trunc (green), RBP-N (violet), the RBP-N/RBP-Trunc dimer (blue) and the full-length RBP (black) without ribose and **(B)** with 0.5 mM ribose. Experiments were performed in 10 mM sodium phosphate, 50 mM sodium chloride, pH 7.8 and the physical and chemical baselines have been subtracted. **(C)** Representative ITC measurement for ribose binding of the RBP-N/RBP-Trunc dimer. Baseline-subtracted raw data are shown at the top while the binding isotherms (blue circles) fitted to a 1:1 model (dotted line) are presented at the bottom. ± at the reported parameters indicate the standard deviation of 3 independent experiments. Titrations were performed at 20 °C in 10 mM sodium phosphate, 50 mM sodium chloride, pH 7.8.

**Table 1.**
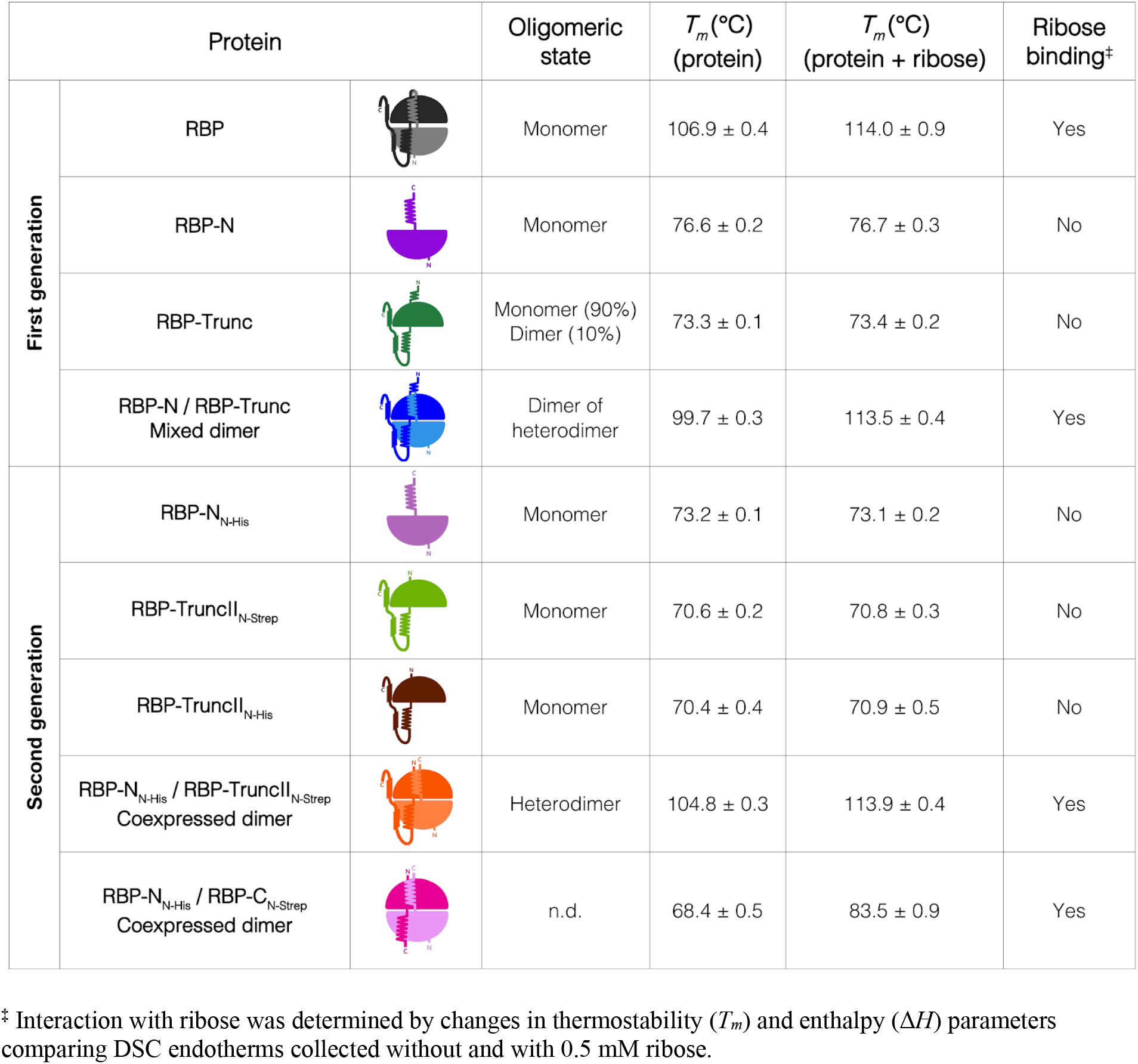
Characterization summary (oligomeric state and thermostability with/without ribose) for the constructs analyzed in this work.

| Protein | | | Oligomeric state | $T_m$ (°C)<br>(protein) | $T_m$ (°C)<br>(protein + ribose) | Ribose binding<sup>‡</sup> |
| --- | --- | --- | --- | --- | --- | --- |
| First generation | RBP |  | Monomer | 106.9 ± 0.4 | 114.0 ± 0.9 | Yes |
|  | RBP-N |  | Monomer | 76.6 ± 0.2 | 76.7 ± 0.3 | No |
|  | RBP-Trunc |  | Monomer (90%)<br>Dimer (10%) | 73.3 ± 0.1 | 73.4 ± 0.2 | No |
|  | RBP-N / RBP-Trunc<br>Mixed dimer |  | Dimer of heterodimer | 99.7 ± 0.3 | 113.5 ± 0.4 | Yes |
| Second generation | RBP-N<sub>N-His</sub> |  | Monomer | 73.2 ± 0.1 | 73.1 ± 0.2 | No |
|  | RBP-TruncII<sub>N-Strep</sub> |  | Monomer | 70.6 ± 0.2 | 70.8 ± 0.3 | No |
|  | RBP-TruncII<sub>N-His</sub> |  | Monomer | 70.4 ± 0.4 | 70.9 ± 0.5 | No |
|  | RBP-N<sub>N-His</sub> / RBP-TruncII<sub>N-Strep</sub><br>Coexpressed dimer |  | Heterodimer | 104.8 ± 0.3 | 113.9 ± 0.4 | Yes |
|  | RBP-N<sub>N-His</sub> / RBP-C<sub>N-Strep</sub><br>Coexpressed dimer |  | n.d. | 68.4 ± 0.5 | 83.5 ± 0.9 | Yes |

<sup>‡</sup> Interaction with ribose was determined by changes in thermostability ( $T_m$ ) and enthalpy ( $\Delta H$ ) parameters comparing DSC endotherms collected without and with 0.5 mM ribose.

The same tendency is observed when comparing the changes in Δ*H* of the individual and mixed constructs (Table S3), with a considerable increase of 240 kcal mol^-1^ in the unfolding enthalpy, which is significantly higher than only the sum of the individual halves (115 kcal mol^-1^). These differences indicate that more accessible surface area is exposed upon unfolding, which is most likely due to the formation of an extensive interface and interdomain interactions important for protein stability and function as present in RBP, confirming the interaction between RBP-N and RBP-Trunc. These results exhibit a similar behavior as observed in the lysine-arginine-ornithine (LAO) binding protein (Vergara, 2023) but differ from those of a previous study of the type-II PBP protein HisJ (Chu, 2013) where the isolated lobes do not interact with each other in the presence or absence of histidine, suggesting that in HisJ only one lobe is important for ligand binding and the other is considered to play a supporting role in the dynamics of binding and in protein stability.

The differences in *T_m_* and Δ*H* of the native proteins and the mixed dimer can be explained by the carry-over of ribose from the purification. It is notoriously hard to remove bound ligands from the expression medium when purifying solute binding proteins that have a high affinity for their ligands (SGC, 2008). Due to its high stability and irreversible thermal unfolding, RBP resisted all attempts of refolding, making purification of a sample removed of all residual ribose not possible, and for this reason always some ribose was carried-over in the purified RBP, increasing the measured *T_m_* and Δ*H* by a ligand stabilization mechanism. Since the individual halves of RBP do not show any binding of ribose (Fig. S3), carry-over is not expected to occur during purification, therefore no additional stabilizing effect of ribose binding is expected.

Next, we determined the crystal structure of the RBP-N/RBP-Trunc dimer (Fig. 4 and Table S4). The two halves indeed reconstitute the canonical RBP fold with high structural similarity, showing a Cɑ-RMSD of 0.41 Å of the dimer to the previously reported structure of unliganded RBP (PDB ID: 2FN9), confirming the aforementioned spectroscopic and calorimetric results. The dimer displays the same opening and twisting angle as the paternal protein, an important indicator of a native-like configuration of the dimer. The asymmetric unit of the crystal structure shows a dimer of RBP-N/RBP-Trunc dimers (Fig. S5), which is in agreement to the oligomeric state observed in SEC-MALS experiments (Fig. 2); however, further analysis is needed to determine the precise conformation of the dimer in solution. The observed dimer interface in the asymmetric unit is mostly related to the interaction of C-terminal residues of RBP-Trunc located in the hinge region and their corresponding ones in the crystallography mates, ruling out the possibility that dimerization results from the extra elements left out in RPB-Trunc. Finally, a closer look at the side-chains involved in ribose binding reveals an almost identical orientation compared to the unliganded state of the native RBP, suggesting the correct formation of the preformed binding site (Fig. 4B). Since all these results showed that the separately purified RBP halves can reassemble the structural conformation of full-length RBP *in vitro*, we next wanted to determine whether this RBP-N/RBP-Trunc dimer is also a functional RBP protein.

**Figure 4.**
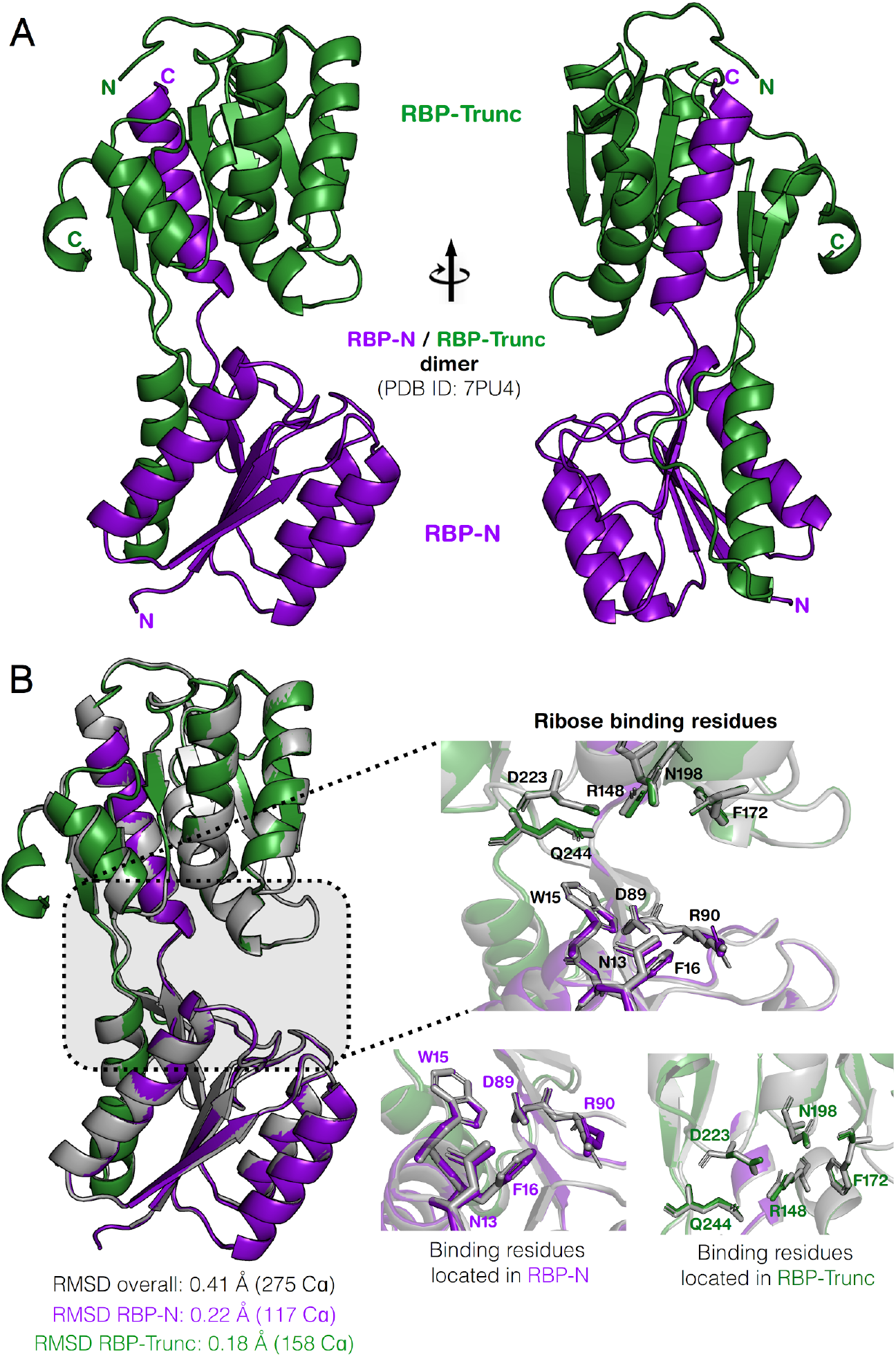
Crystal structure of the RBP-N/RBP-Trunc dimer in unliganded conformation. **(A)** Cartoon representation of RBP-N (violet) and RBP-Trunc (green) dimer (PDB ID: 7PU4) forming a native-like conformation as full-length RBP. **(B)** Structural comparison of RBP-N/RBP-Trunc dimer and RBP (PDB ID: 2FN9; grey). RMSD values are reported for the entire dimer, and halves RBP-N and RBP-Trunc. Inset shows the ribose binding residues in the full structure (top) and separated in each half (bottom); numbering is based on the RBP sequence.

### The reassembled heterodimer binds ribose with a comparable affinity to full-length RBP

The structural similarity of the dimer with the full-length RBP suggests that also the ribose binding function might be reconstituted. To investigate this, we first analyzed by DSC if ribose binding increases protein thermostability. Specific protein-ligand interaction commonly causes an increase in protein thermostability, which is due to the coupling between binding and unfolding processes under thermodynamic equilibrium (Cooper, 2000; Privalov, 1979). While the isolated RBP-N and RBP-Trunc do not show any sign of stabilization by ligand binding (Fig. S3), an increase in *T_m_* can be observed upon addition of 0.5 mM ribose (Fig. 3B) to the RBP-N/RBP-Trunc dimer, with the amplitude of the absorbed heat changes being dependent on ligand concentration. The *T_m_* of the ligand-bound RBP-N/RBP-Trunc dimer increases by almost 14 °C from 99.7 to 113.5 °C, comparable to the stabilization of ligand-bound RBP by around 7 °C to 114.0 °C (Fig. S3 and Table S3) and similar to the one observed in other PBPs when binding their respective high-affinity ligands (Kreimer, 2003; Ganesh, 1997; Fukada, 1983).

In addition, an increase of 129 kcal mol^-1^ was observed in the unfolding Δ*H* for the ligand-bound RBP-N/RBP-Trunc dimer in comparison to the unbound form, deducing that large-scale rearrangements in the solvent-exposed surface in the dimer accompanies ligand binding, thereby confirming a functional protein that behaves similar to full-length RBP. The greater amount of thermostabilization in the dimer in comparison to RBP can again be explained by residual ribose carried over in the purification of RBP already stabilizing the protein. However, at the same concentration of ribose the level of stabilization of the dimer is almost identical to that of RBP, with the dimer displaying a native-like thermostability. Interestingly, the significant increase in stability can also be observed when adding ribose to a non-native SDS-PAGE. At concentration of 1 mM ribose or higher, a dimer (and higher oligomers) can be detected, indicating that the addition of SDS and the subsequent heating to 99 °C is not enough to dissociate the ribose-bound stabilized dimer (Fig. S4).

Additionally to DSC analysis, ribose binding of the RBP-N/RBP-Trunc dimer was determined by ITC. Ribose-binding isotherms (Fig. 3C) showed a sigmoidal profile with the ribose binding constant (*K*_D_ = 6.8 ± 0.7 µM) in a concentration range comparable to other previously studied solute binding proteins (Schreier, 2009), implying that the binding of ribose can be regained after *in vitro* mixing the previously dissected RBP halves. In fact, ligand affinity is not significantly affected by the assembly. Now the question remained, whether this reassembled functional dimer can also be formed *in vivo* upon co-expression of both halves.

### RBP halves form a functional dimer when co-expressed in E. coli

To investigate whether the dimer of RBP-N and RBP-Trunc already forms during the expression in *E. coli*, a second generation of constructs was created (Table S1). To ensure that at least one plasmid copy of each construct stays in each cell, the coding sequences were assembled in a vector imparting resistance to either ampicillin or kanamycin, respectively. Since there was no control of expression levels and we wanted to only obtain homodimer in the subsequent purification, we opted for adding two different affinity tags to each construct (Fig. 5A). The resulting constructs are RBP-N_N-His_ and RBP-TruncII_N-Strep_ (Table S1) with affinity labels located at the N-terminus. By utilizing a three-step purification approach using the different affinity tags on each protein half and a subsequent SEC step for polishing, we can assure that only already formed dimer is retained as confirmed by the SDS-PAGE showing a band at the corresponding sizes of both RBP-N_N-His_ and RBP-TruncII_N-Strep_ and thermal resistance upon addition of ribose (Fig. 5B). Similar to the behavior of the 1^st^ generation constructs, the far-UV CD and fluorescence spectra showed a reconstitution of characteristics almost identical to the native RBP (Fig. S2B and Fig. S6A). The molecular weight determined by SEC-MALS also corresponds to the dimer (expected mass: 36.8 kDa / determined mass: 37.3 kDa), with no higher oligomers present (Fig. 5C and Table S2).

**Figure 5.**
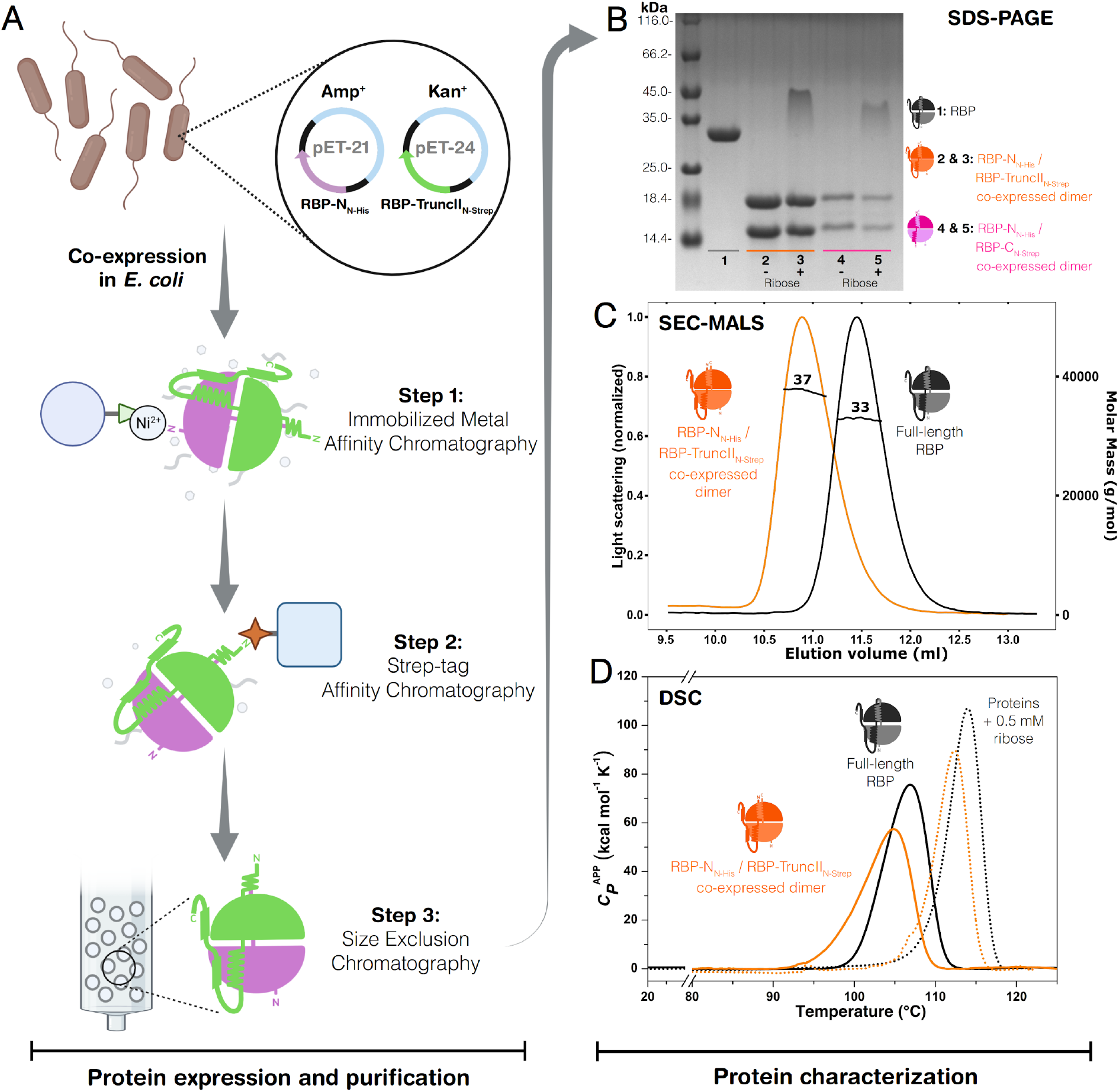
Co-expression in *Escherichia coli* and characterization of the second-generation dimers. **(A)** Schematic workflow of the co-expression beginning with the transformation of *E. coli* with the two plasmids carrying RBP-N_N-His_ and RBP-TruncII_N-Strep_. Subsequent alternating affinity chromatographies utilizing two different tags assure purification of only the RBP-N_N-His_/RBP-TruncII_N-Strep_ dimer, followed by a final size exclusion step. **(B)** SDS-PAGE showing the co-purified dimers. RBP (lane 1), co-expressed RBP-N_N-His_/RBP-TruncII_N-Strep_ dimer without ribose (lane2) and with 0.5 mM ribose (lane3), co-expressed RBP-N_N-His_/RBP-C_N-Strep_ dimer without ribose (lane 4) and with 0.5 mM ribose (lane 5). **(C)** SEC-MALS measurements of RBP-N_N-His_/RBP-TruncII_N-Strep_ dimer (orange line) in comparison with full-length RBP (black line). Numbers indicate the determined experimental molecular weight. **(D)** DSC endotherms of RBP-N_N-His_/RBP-TruncII_N-Strep_ dimer and RBP in absence (continuous lines) and presence of 0.5 mM ribose (dotted lines).

Similar to the mixed RBP-N/RBP-Trunc dimer, the co-expressed and co-purified dimer shows an increase in thermostability in the presence of ribose (Fig. 5D and Table 1), indicating a functional dimer. The *T_m_* of RBP-N_N-His_/RBP-TruncII_N-Strep_ increases by 9.1 °C (from 104.8 °C to 113.9 °C) after addition of 0.5 mM ribose, showing a similar trend of stabilization as the full-length RBP (Fig. S3 and Table S3) and also to the 1^st^ generation of halves. In view of the dimer being formed in the cells during co-expression, the same behavior of carrying over residual ribose from *E. coli* is expected to increase the measurable *T_m_* of the RBP-N_N-His_/RBP-TruncII_N-Strep_ dimer.

Since the formation of the dimer appears to stabilize the individual protein halves and yields properties almost identical to full-length RBP, we set out to retry the expression of the previously insolubly expressing RBP-C in the hopes that the co-expression and formation of the dimer *in-cell* could rescue the protein. The 2^nd^ generation RBP-C_N-Strep_ was purified along RBP-N_N-His_ analogously to the previous co-expression assay (Fig. 5A). Interestingly, we were able to obtain a small amount of purified dimer after the affinity chromatography and subsequent SEC (Fig. 5B), with a dimer band still being visible in the SDS-PAGE after addition of ribose, which indicates retention of binding function. This characteristic is confirmed by DSC measurements of the dimer with and without ribose (Fig. S7). While the overall transition is massively decreased for the unbound proteins (*T_m_* = 68.4 °C for RBP-N_N-His_/RBP-C_N-His_ dimer vs. *T_m_* = 106.9 °C for full-length RBP), the strong stabilization after addition of ribose is still observed (15.1 °C of *T_m_* increase to 83.5 °C). The total shift is comparable with that in RBP, albeit with some fraction of the protein still appearing to be in a ligand-free state (Fig. S7), indicating a possible reduction in ribose binding affinity or different populations of the purified dimer. The ability of RBP-N_N-His_ to recover not just the soluble expression of RBP-C_N-Strep_ via the formation of the dimer, but also the dimer to retain its function, showcases the inherent versatility of this fold and gives insights into its evolution.

## Conclusions

### Implications for the evolution of the PBP fold and protein engineering approaches

The data presented here shows how a modern PBP can be disassembled into its two lobes, and how when they are combined *in vitro* or *in vivo* the formed dimer is able to perform its original function. The individual parts readily assemble to form a dimer, not just when mixing the individually purified lobes, but also within the cell upon co-expression. While the N- and C-terminal lobes appear to be stable and well-behaved proteins on their own, formation of the dimer almost completely restores the characteristics of the full-length RBP, confirming the importance of interdomain interactions on the evolution, stability and function of the PBP fold, similarly to what has been reported for other multidomain proteins (Han, 2007; Vogel, 2004).

Analysis of the stability and binding abilities indicate native-like properties, and the crystal structure of the heterodimer being nearly identical to that reported for RBP supports this conclusion. This versatility of the PBP fold can be explained by the inherent malleability of proteins of the flavodoxin-like (and related) folds. Several structures with swapped elements have been reported for flavodoxin-like proteins (e.g. PDBs: 4Q37, 6ER7/6EXR, 3C85; Paithankar, 2019; Farías-Rico, 2014) as well as TIM-barrel proteins (PDB 6QKY; Michalska, 2020), which are also thought to be related to the flavodoxin-like fold (Romero-Romero, 2021). Further, we had previously observed swapped elements in circular-permuted constructs of RBP (PDBs: 7QSP, 7QSQ; Michel, 2023). This tendency of the structural archetype to enable formation of swapped elements could have been an important characteristic promoting the emergence of the ancestral dimer thought to be the progenitor of modern PBPs. While the two halves we describe in this work are derived from an already evolved protein, they could still be seen as a vestige of this ancestral dimer. Interestingly, the crystal structure of a flavodoxin-like fold protein with an identical arrangement of secondary structure elements has been described already, albeit it is unclear whether the observed structure is an artifact of the non-physiological crystallographic conditions (Lewis, 2000).

Since the dimer corresponds to the proposed ancestral dimer in the evolutionary trajectory (Fig. 1A) while still retaining function with native-like properties, this presents new insight into the mechanisms behind such a duplication event. Not only does the orientation of the two lobes create the binding cleft characteristic for PBP-like proteins, but also the general restraints on the movement of the lobes lower the entropic cost of ligand binding. Our findings showcase the feasibility of a functional dimer similar to the proposed ancestral one to also assemble within cells, giving way to the argument that the duplication and fusion of the progenitor flavodoxin-like protein might have happened independent of the gain of function, indicating no evolutive pressure on single domains but on the full-length RBP.

Adopting this approach and expanding it to incorporate a diverse set of functions could also be used for protein engineering purposes. This is traditionally done by inserting a domain for readout into the sequence of an existing PBP, with the optimal placement of the insertion sites being one of the major challenges (Ribeiro, 2019; Tullman, 2016). Further studies will have to show that the retracing of the duplication is applicable for other PBPs as well, but one could imagine its usage in creating modular switch systems not just *in-vitro* but also *in-vivo*.

## Materials and Methods

### Reagents and solutions

Analytical grade chemicals were used for all the experiments. Water was distilled and deionized.

### Identification of the protein halves and sequence analysis

The bioinformatic analysis to trace the sequence similarities between the RBP and flavodoxin-like proteins was done using the HHpred server which is part of the HHsuite (Gabler, 2020) (Fig. S1). The sequence of full-length RBP (UniProt-ID: Q9X053) excluding the extracellular transport signal was run with standard parameters, but disabling secondary structure scoring and increasing the number of maximal hits to 10000 to also obtain sequences with lower probability scores. Based on the alignment of both the other PBP lobes and the hits with the flavodoxin-like proteins, the cutting points were determined at position 30-155 for RBP-N, 142-310 for RBP-Trunc,156-310 for RBP-TruncII and 157-291 for RBP-C (Table S1).

### Cloning and generation of RBP-constructs

The gene fragment for wild type RBP lacking the periplasmic signal sequence as well as the primers used for assembly were provided by Eurofins Genomics. To generate the gene fragments for RBP-N and RBP-Trunc, a polymerase chain-reaction with the corresponding primer was conducted with the full sequence as template. Additionally, a QuikChange® site-directed mutagenesis was performed to obtain the M142A mutation of the full-length RBP to prevent the translation of the truncated protein (henceforth called RBP). The fragment of full length RBP was cloned into empty pET-21 using the *Nde*I/*Xho* I restriction sites. Analogously generated fragments for RBP, RBP-N, and RBP-Trunc were all subsequently cloned using T5 exonuclease-dependent assembly (Xia, 2019). All constructs were verified by sequencing.

Gene synthesis and cloning for the co-expression assay were provided by Biocat. The differently tagged constructs of RBP-TruncII and RBP-N_N-His_ were cloned into pET24-and pET21-vectors, respectively. Individual clones were obtained by transforming *Escherichia coli* BL21 (DE3) cells by adding 50 ng of purified plasmid, heat shock and subsequent plating on agar-plates supplemented with the corresponding antibiotic. To obtain cells carrying the two different plasmids needed for the co-expression assay, 50 ng of each plasmid were added to the *E. coli* BL21 (DE3) cells, heat shocked and then grown on plates containing the two selecting antibiotics.

### Expression and purification of RBP-constructs

The transformant *E. coli* BL21(DE3) were grown in *Terrific broth* media (TB) at 37 °C to an OD_600_ of 1.2 in the presence of the corresponding antibiotics (ampicillin 100 µg mL^-1^; kanamycin 50 µg mL^-1^). Protein expression was induced by the addition of Isopropyl-β-thiogalactopyranoside to a concentration of 1 mM and a total time of 18 h at 20 °C. Cells were harvested via centrifugation (5000 × G, 15 min), resuspended in the corresponding binding buffer (20 mL g^-1^ wet weight), lysed by sonication and subsequently centrifuged to remove remaining cell debris (40000 × G, 1 h). The cleared lysate was filtered through a 0.22 µm filter previous to the affinity column step.

For the constructs carrying a hexahistidine affinity tag, Immobilized Metal Ion Chromatography (IMAC) was performed on a Cytiva HisTrap 5 mL column equilibrated with buffer (20 mM MOPS, 500 mM sodium chloride, 10 mM imidazole, pH 7.8). Elution was performed with a step of IMAC-Elution-Buffer (20 mM MOPS, 500 mM sodium chloride, 600 mM imidazole, pH 7.8) at 40%, and fractions corresponding to the eluted protein pooled and concentrated to a volume suitable for the size exclusion chromatography step.

Strep-Tactin affinity chromatography was used for constructs with a StrepII-Tag, which were loaded onto a Cytiva StrepTrap HP 5mL column equilibrated with Strep-Trap binding Buffer (100 mM Tris-HCl, 150 mM sodium chloride, 1 mM EDTA, pH 7.8) and eluted with Strep-Trap elution Buffer (100 mM Tris-HCl, 150 mM sodium chloride, 1 mM EDTA, 2.5 mM Desthiobiotin, pH 7.8), pooled and concentrated analogous to the IMAC purification. To facilitate purification of the individual constructs, the Strep-Tag of RBP-Trunc_N-Strep_ was switched to a His_6_-Tag, creating RBP-Trunc_N-His_.

For the purification of the co-expressed constructs, to assure survival of cells carrying only the two plasmids, the LB medium used for the production was supplemented with both Ampicillin and Kanamycin (100 µg mL^-1^ and 50 µg mL^-1^, respectively). Cell lysis was performed as with the individual constructs, and the lysate first loaded on the HisTrap column. The eluted fractions corresponding to the tagged protein were pooled and applied onto a StrepTrap column. Similarly, eluted fractions were pooled and concentrated to a volume suitable for application onto the Superdex column.

Size exclusion chromatography was performed as final purification step for all constructs on a Cytiva Superdex 26/600 75 pg with an isocratic elution using buffer 10 mM sodium phosphate, 50 mM sodium chloride, pH 7.8. Fractions consistent with the proteins of interest were analyzed by SDS-PAGE, pooled, flash frozen in liquid nitrogen, and stored at -20 °C until further analysis.

### Far-UV Circular Dichroism (CD)

Far-UV Circular Dichroism (CD) measurements were performed at 20 °C in buffer 10 mM sodium phosphate, 50 mM sodium chloride, pH 7.8 in a Jasco J-710 spectropolarimeter equipped with a Peltier device to control temperature (PTC-348 WI). Spectra were collected using 5 µM protein concentration for RBP and the dimers, and 10 µM for the other constructs in a 2 mm cuvette, 195-260 nm wavelength range, and 1 nm bandwidth. After buffer subtraction, raw data were converted to mean residue molar ellipticity ([*Θ*]) with [*Θ*] = *Θ / l C N_r_*, where *Θ* is the ellipticity signal in millidegrees, *l* is the cell path in mm, *C* is the molar protein concentration, and *N*_r_ is the number of amino acids per protein (Greenfield, 2007).

### Intrinsic Fluorescence (IF)

Intrinsic fluorescence (IF) spectra were collected on a Jasco FP-6500 spectrofluorometer coupled with a water bath (Julabo MB) to control the temperature. Experiments were performed at 20 °C in buffer 10 mM sodium phosphate, 50 mM sodium chloride, pH 7.8 and 5 µM protein concentration for RBP and dimers, and 5 µM for the other proteins, with 280 nm as excitation wavelength, 300-500 nm as emission wavelength, and 1 nm bandwidth. Raw signal was normalized for protein concentration.

### Analytical Size Exclusion Chromatography coupled to Multi Angle Light Scattering (SEC-MALS)

Analytical Size Exclusion Chromatography measurements were performed coupled to a miniDAWN Multi Angle Light Scattering (MALS) detector and an Optilab refractometer (Wyatt Technology). Samples previously centrifuged and filtered were run in a Superdex 75 Increase 10/300 GL column connected to an Äkta Pure System (GE Healthcare Life Sciences) equilibrated with buffer 10 mM sodium phosphate, 50 mM sodium chloride, 0.02% sodium azide, pH 7.8. Experiments were conducted at room temperature with a protein concentration of 1 mg mL^-1^ and 0.8 mL min^-1^ flow rate. For the samples containing ribose, 0.5 mM of ribose was premixed with protein at 1 mg mL^-1^. Reproducibility during all SEC-MALS measurements was tested by running a BSA standard at 2 mg mL^-1^ at the beginning and end of all experiments, which resulted in identical data. Determination of weight averaged molar mass was performed by using the Zimm-Equation with the differential refractive index signal as source for the concentration calculations (refractive index increment dn/dc set to 0.185). Data collection and analysis were done using ASTRA v.7.3.2 software (Wyatt Technology).

### Crystallization and three-dimensional structure determination

For setting up crystallization assays, protein at 0.5 mM concentration was dialyzed against 20 mM Tris-HCl, 300 mM sodium chloride, pH 7.8. For RBP-N / RBP-Trunc dimer, 0.5 mM equimolar ratio of each protein was used as initial concentration. Screening plates were set up by a sitting-drop vapor diffusion method using JCSG Core I-IV (Qiagen), PEG Suite I-II (Qiagen), and Additive Screen kits (Hampton Research) in 96 well Intelli plates (Art Robbins Instruments). Plates with 0.8 µL drops in a 1:1, 1:2, and 2:1 protein:mother liquor drop ratio were set up with a nano dispensing crystallization Phoenix robot (Art Robbins Instruments) and stored at 20 °C in a hotel-based crystal imaging system RockImager RI 1000 (Formulatrix). RBP-N / RBP-Trunc dimer crystals with successful diffraction data were found in 100 mM HEPES pH 7.5, 15 % (w/v) PEG 20000 and a drop ratio 1:1. Data were collected at Berlin Electron Storage Ring Society for Synchrotron Radiation beamline 14.2 (BESSY 14.2) operated by the Helmholtz-Zentrum Berlin using the mxCuBE beamline-control software (Gabadinho, 2010). Measurements at 100 K were performed in a single-wavelength mode at 0.9184 Å with a PILATUS3S 2M detector (HZB, 2016) in fine-slicing mode (0.1° wedges). Diffraction images were processed with X-ray Detector Software (XDS) and XDSAPP v3.0 (Kabsch, 2010; Sparta, 2016). Phasing was performed by molecular replacement with PHASER in the PHENIX software suite v.1.19.2 (Liebschner, 2019) using the edited pdb file corresponding to the RBP-N and RBP-Trunc halves from *Thermotoga maritima* RBP (PDB 2FN9). Data refinement was carried out with phenix.refine (Adams, 2010) and iterative manual model building/improvement in COOT v.0.9 (Emsley, 2010). Coordinates and structure factors were validated and deposited in the PDB database https://www.rcsb.org/ (Berman, 2002) with the accession code: 7PU4. Figures were created with PyMOL Molecular Graphics System v.2.3.0 (Schrodinger, LLC).

### Differential Scanning Calorimetry (DSC)

Differential Scanning Calorimetry endotherms were collected using a VP-Capillary DSC instrument (Malvern Panalytical) with a temperature range of 10-130 °C and 1.5 °C min^-1^ scan rate. Protein samples were prepared at 50 µM after exhaustive dialysis in buffer 10 mM sodium phosphate, 50 mM sodium chloride, pH 7.8 and proper degassing. Instrument equilibration was performed by collecting at least 2 buffer-buffer scans before each protein-buffer experiment. Calorimetric reversibility was tested by collecting two consecutive endotherms and calculating the recovery area percentage from the second and first scan, resulting in irreversible thermal-unfolding transitions for all the constructs reported in the present study. Thermodynamic parameters (*T*_m_ and Δ*H*) were calculated after subtracting physical (buffer-buffer scan) and chemical baselines (heat capacity effects) from each protein-buffer scan. Thermostabilization by protein-protein interaction (dimer formation) was determined by changes in *T*_m_ and Δ*H* when two different proteins were combined in equimolar concentration. DSC experiments in presence of ribose were performed at 50 µM protein concentration and 0.5 mM ribose premixed in the same working buffer before the heating cycles. Buffer-buffer scans were collected containing the same amount of ribose as protein/ribose-buffer experiments and subtracted as indicated. Ribose stability at high temperatures was tested and no endotherm distortions were observed in the concentration and temperature ranges assayed. Origin v.7.0 (OriginLab Corporation) with MicroCal software was used for data analysis.

### Isothermal Titration Calorimetry (ITC)

Binding assays followed by Isothermal Titration Calorimetry (ITC) were performed using a TA Nano ITC low volume device (TA Instruments). Titrations were obtained at 20 °C in buffer 10 mM sodium phosphate, 50 mM sodium chloride, pH 7.8 and 100 µM of protein concentration, which was exhaustively dialyzed against the working buffer. Ribose solution was prepared in the same working buffer to minimize dilution heats and was loaded in the syringe at 0.8 mM concentration. Protein and ligand solutions were degassed with a vacuum pump for 90 mins before carrying out the experiments, and concentrations were optimized in order to reach *c* values higher than 10. Independent triplicates of ITC experiments were performed with 25 injections of 2 µL volume, spacing of 350 s between injections, and stirring at 300 rpm. Dilution heats were subtracted from the heat associated with each injection to get accurate parameters. Baseline and integration intervals were carefully checked to avoid experiment distortions. Binding constant (*K*_D_), enthalpy change (Δ*H*), and binding stoichiometry (*n*) were determined by nonlinear fitting of normalized data assuming a 1:1 binding model and using TA ITC software. All titration replicates fulfilled the characteristics for an accurate parameter determination that have been analyzed by experimental and simulation data (Turnbull, 2003).

## Abbreviations

CD: Circular Dichroism
DSC: Differential Scanning Calorimetry
IF: Intrinsic Fluorescence
ITC: Isothermal Titration Calorimetry
MALS: Multi Angle Light Scattering
PBP: periplasmic binding protein
RBP: ribose-binding protein
SEC: size exclusion chromatography
*T_m_*: midpoint of thermal unfolding
Δ*H*: change in enthalpy

## Acknowledgments

We acknowledge allocation of synchrotron beamtime and financial support by HZB and thank the beamline staff at BESSY for support. We thank Saacnicteh Toledo-Patiño for scientific discussions and input in the early stages of the project, Sabrina Wischt and Sooruban Shanmugaratnam for their competent technical support, and all the members of the Höcker Lab for their constructive suggestions to improve the research.

## Funding

This work was supported by the European Research Council (ERC Consolidator Grant 647548 ‘Protein Lego’ to B.H.), the VolkswagenStiftung (grant 94747 to B.H.), and by a fellowship from the Alexander von Humboldt and Bayer Science & Education Foundation (Humboldt-Bayer Research Fellowship for Postdoctoral Researchers to S.R.R.).

## Competing interests

The authors declare that they have no conflicts of interest with the contents of this article.

## Author Contributions

F.M., S.R.R., B.H. designed the research, F.M., S.R.R. purified the different constructs, F.M. collected and analyzed CD, IF, and SEC-MALS data, S.R.R. performed DSC and ITC experiments, F.M., S.R.R. crystallized and solved three-dimensional structure, F.M., S.R.R., B.H. wrote the manuscript.

## Data and materials availability

All data to support the conclusions of this manuscript are included in the main text and supplementary materials. Coordinates and structure factors have been deposited to the Protein Data Bank (PDB) with accession code: 7PU4 (RBP-N / RBP-Trunc dimer).

## Supplementary Materials

This article contains supplementary material that includes: supplementary figures 1-5 and supplementary tables 1-4.

## Supplementary Information

### Supplementary figures and tables

#### List of Supplementary Figures

**Figure S1.**
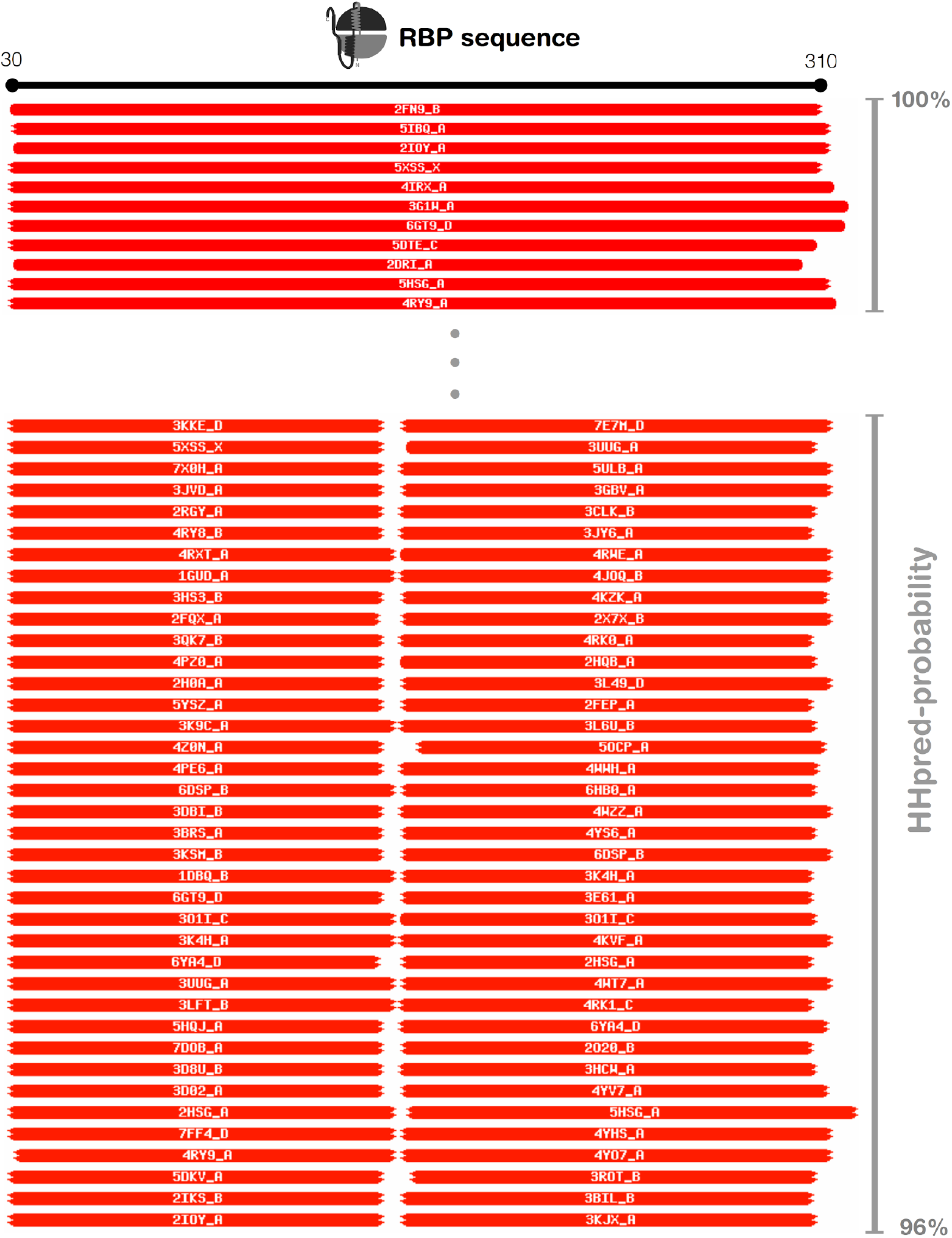
Representative HHpred results for the RBP sequence. Visualization of the HHpred output showing the query sequence as a black bar. The database matches are shown as red horizontal bars underneath with their respective identifiers. Bar length is indicating its coverage with respect to the query and is colored according to its significance (red as very significant to orange, yellow, green and cyan as less significant). Top and longer bars show the alignment of other full-length PBPs on the query sequence while bottom and shorter bars indicate the alignment of the individual lobes. On the right is the HHpred probability shown for the presented sequence range. Numbering has been adapted to be consistent with uniprot entry Q9X053.

**Figure S2.**
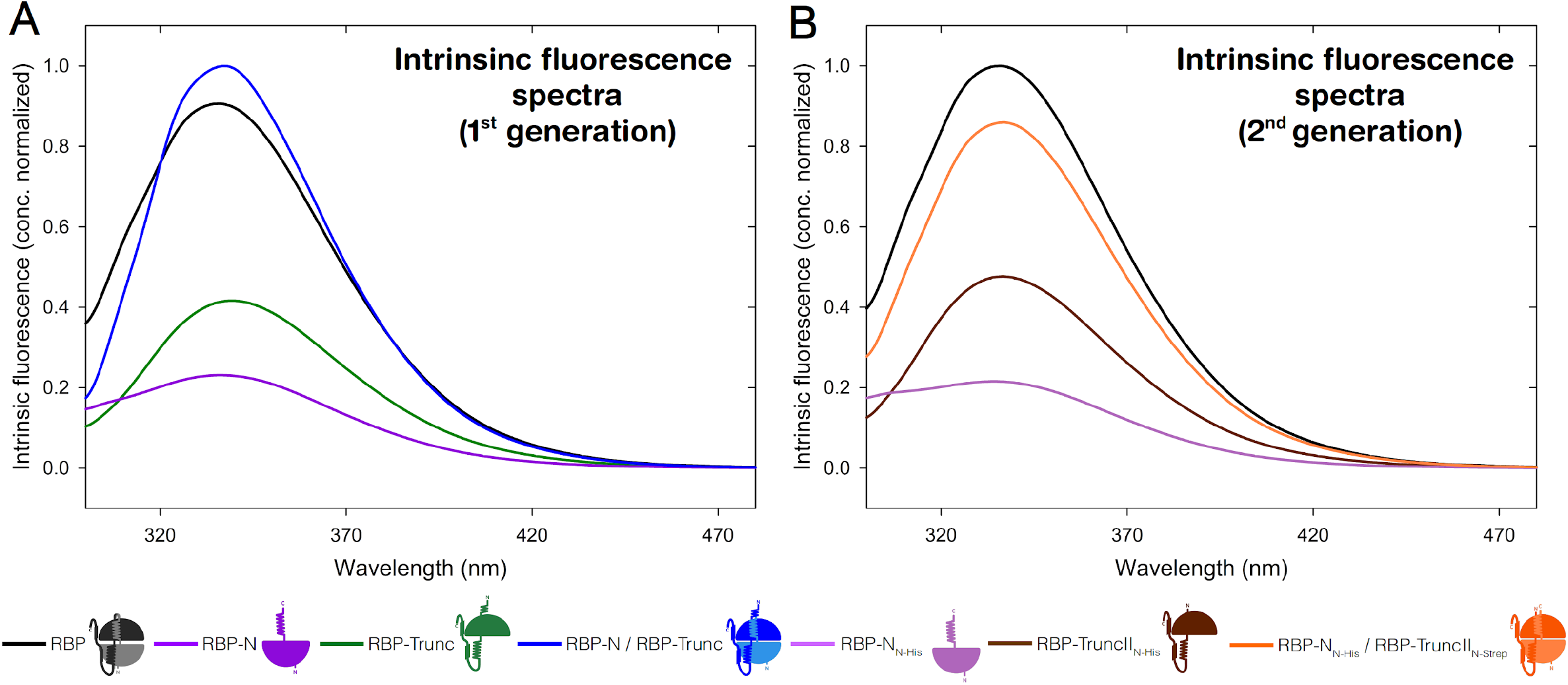
Intrinsic fluorescence measurements of the first- and second-generation constructs. Fluorescence spectra measured from 300-480 nm at an excitation wavelength of 280 nm of RBP (black), RBP-N (violet), RBP-Trunc (green) and the mixed RBP-N/RBP-Trunc dimer (**A**) and RBP-N_N-His_ (red), RBP-TruncII_N-His_ (brown) and the co-expressed RBP-N_N-His_/RBP-TruncII_N-Strep_ dimer (**B**) in 10 mM sodium phosphate, 50 mM sodium chloride, pH 7.8. Signal was normalized by protein concentration.

**Figure S3.**
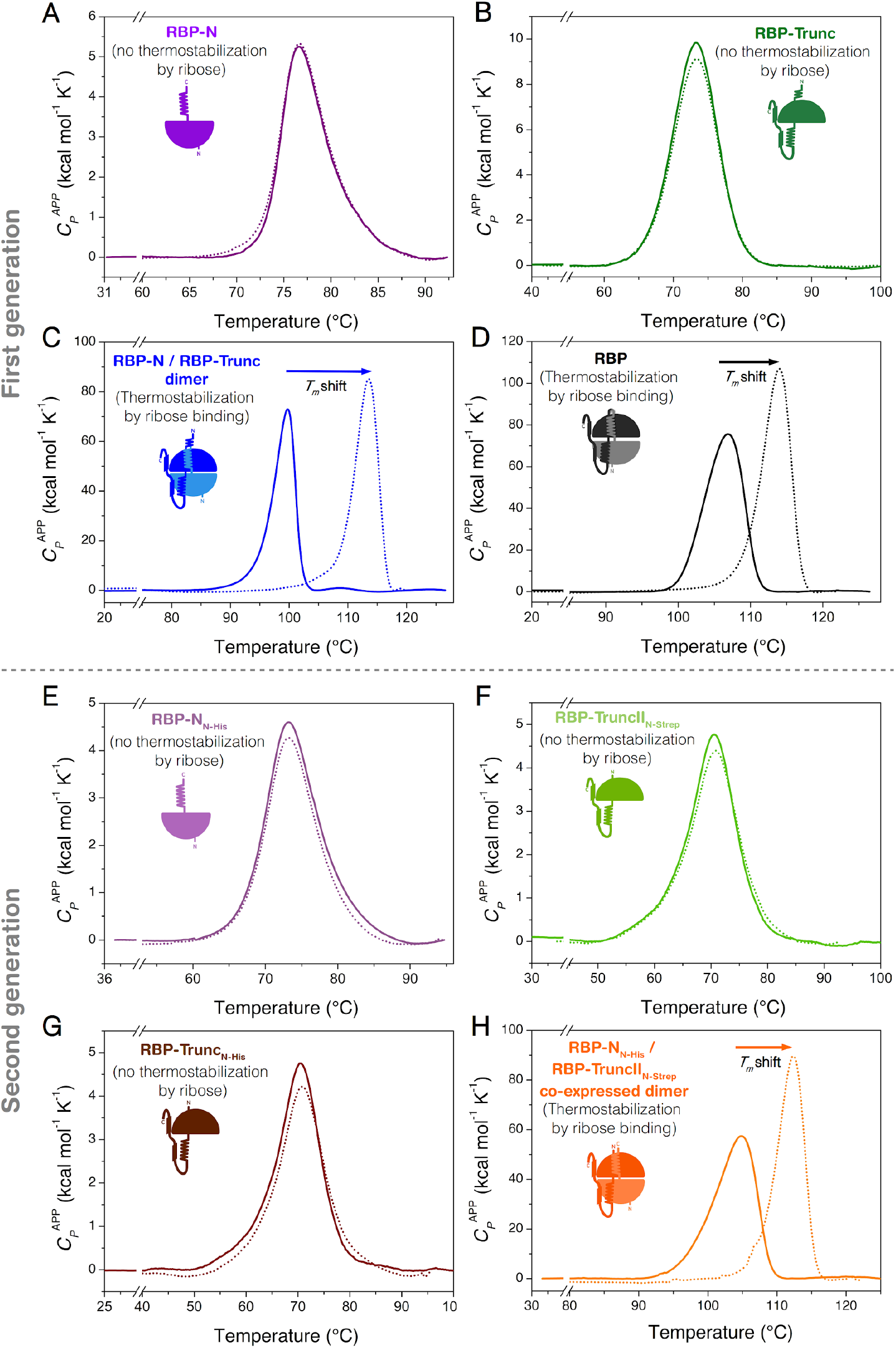
DSC experiments for the first- and second-generation constructs. DSC endotherms at 1.5 °C min^-1^ without ribose (solid lines) and with 0.5 mM ribose (dotted lines) of **(A)** RBP-N (violet), **(B)** RBP-Trunc (green), **(C)** RBP-N/RBP-Trunc dimer, **(D)** full-length RBP (black), **(E)** RBP-N_N-His_ (light purple), **(F)** RBP-TruncII_N-Strep_ (light green), **(G)** RBP-TruncII_N-His_ (brown), and **(H)** co-expressed RBP-N_N-His_/RBP-TruncII_N-Strep_ dimer (orange). Experiments were performed in 10 mM sodium phosphate, 50 mM sodium chloride, pH 7.8 and the physical and chemical baselines have been subtracted.

**Figure S4.**
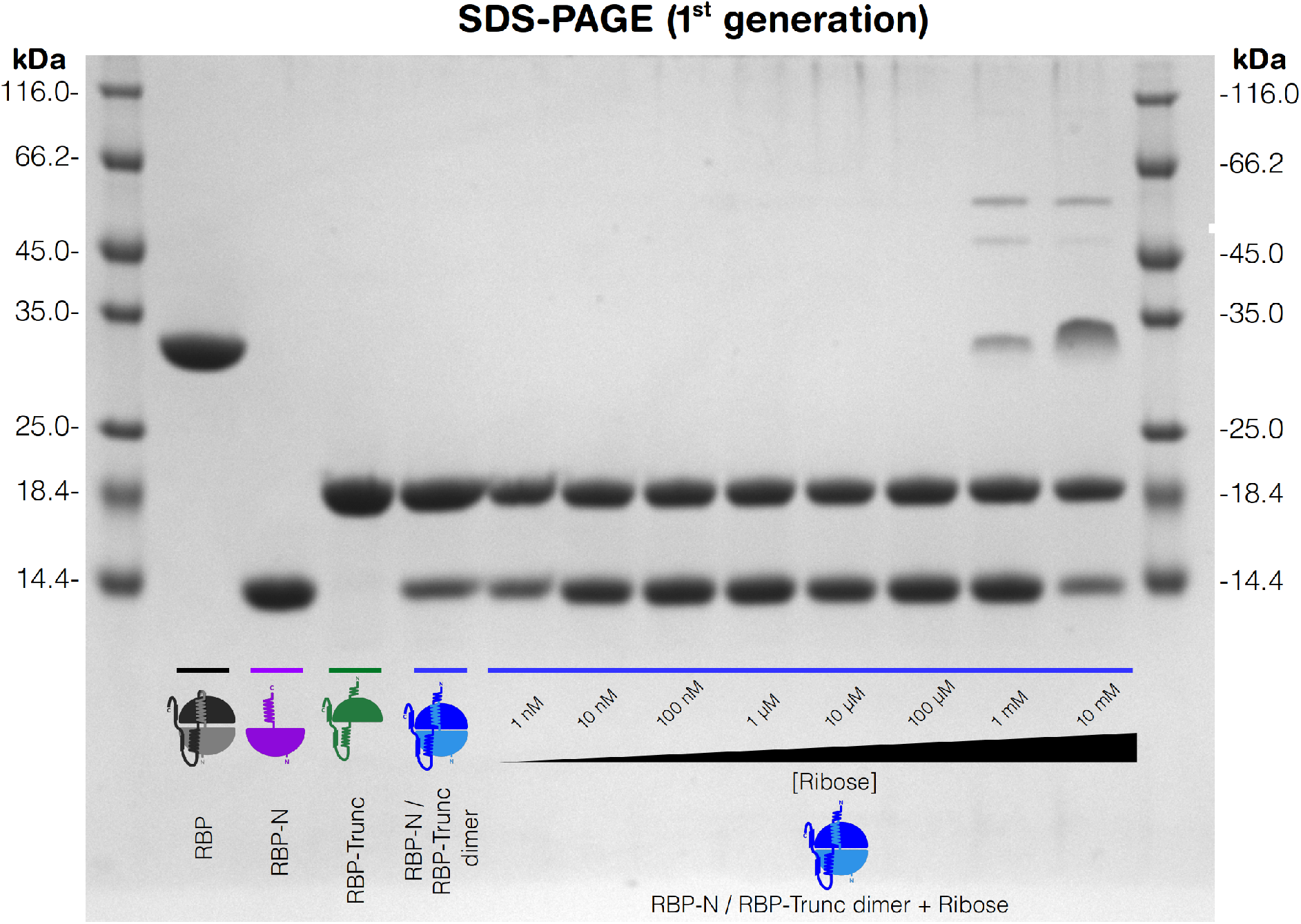
SDS-PAGE of RBP, the individual first-generation halves, and the mixed dimer. Purified RBP, RBP-N, RBP-Trunc and RBP-N/RBP-Trunc dimer (lane 2-5 respectively) show single proteins at the expected molecular weight without major contaminants. Addition of ribose to the dimer appears to stabilize the complex to a degree where it becomes resistant to dissociation in the SDS loading buffer and subsequent heating as indicated by the presence of higher oligomer bands in the presence of ≥1 mM [ribose] (lanes 6-13). Molecular weight has been estimated as indicated by the addition of the molecular weight standard (lane 1 and 14, weights annotated).

**Figure S5.**
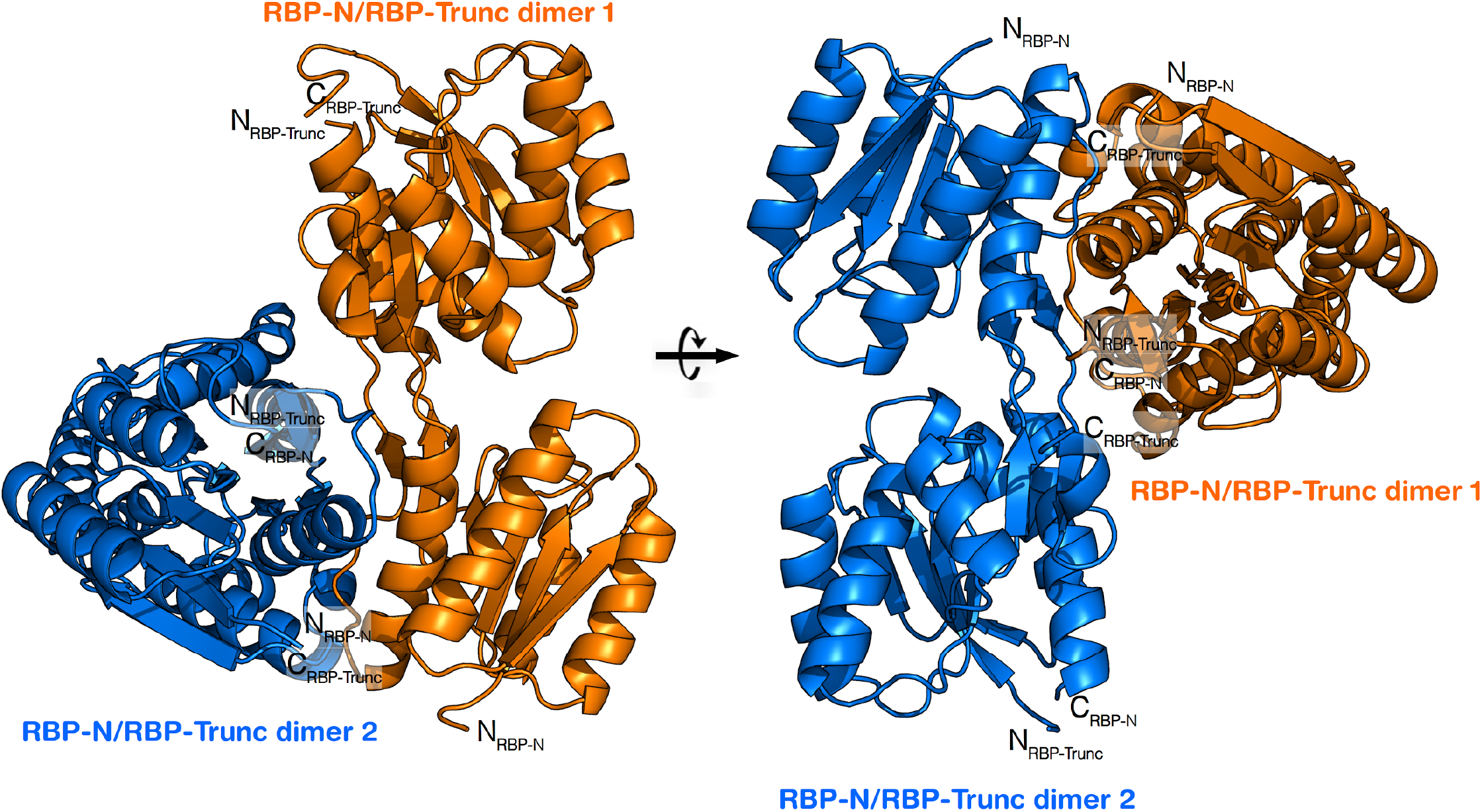
Crystallographic dimer formed by the asymmetric-unit mate of RBP-N/RBP-Trunc crystal structure. Each heterodimer is indicated in orange and blue.

**Figure S6.**
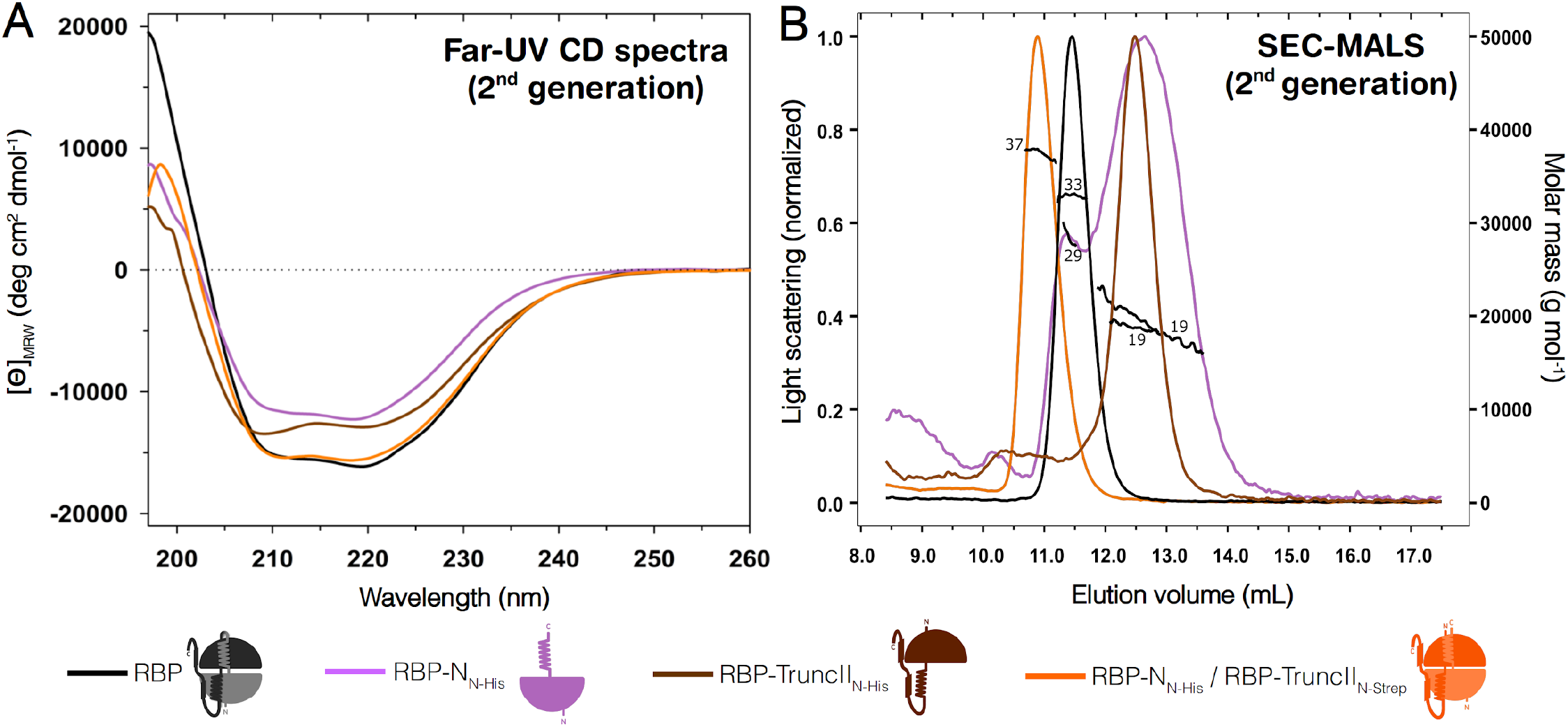
Biophysical characterization of the second-generation constructs. (A) Far-UV CD spectra of RBP (black), RBP-N_N-His_ (light purple), RBP-TruncII_N-His_ (brown) and the coexpressed RBP-N_N-His_/RBP-TruncII_N-Strep_ dimer (orange) in 10 mM sodium phosphate, 50 mM sodium chloride, pH 7.8. **(B)** SEC-MALS measurements in 10 mM sodium phosphate, 50 mM sodium chloride, 0.02% sodium azide, pH 7.8. Numbers indicate the determined molecular weight after data analysis. Values derived from the experiments are reported in Supplementary Table S2.

**Figure S7.**
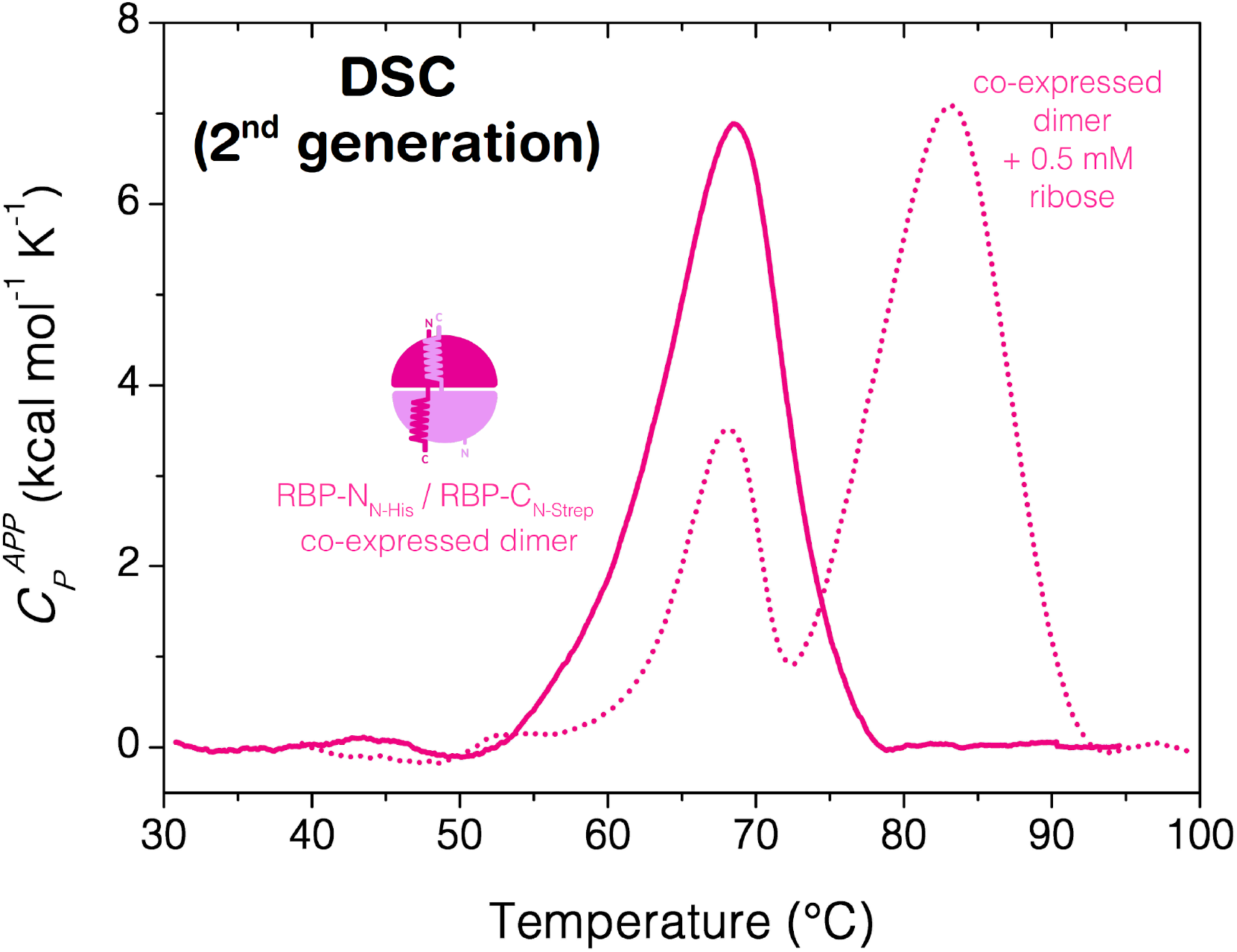
DSC endotherms for the coexpressed RBP-N_N-His_/RBP-C_N-Strep_ dimer. DSC experiments were collected at 1.5 °C min^-1^ without ribose (solid lines) and with 0.5 mM ribose (dotted lines) in 10 mM sodium phosphate, 50 mM sodium chloride, pH 7.8. Physical and chemical baselines were subtracted.

#### List of Supplementary Tables

**Table S1.**
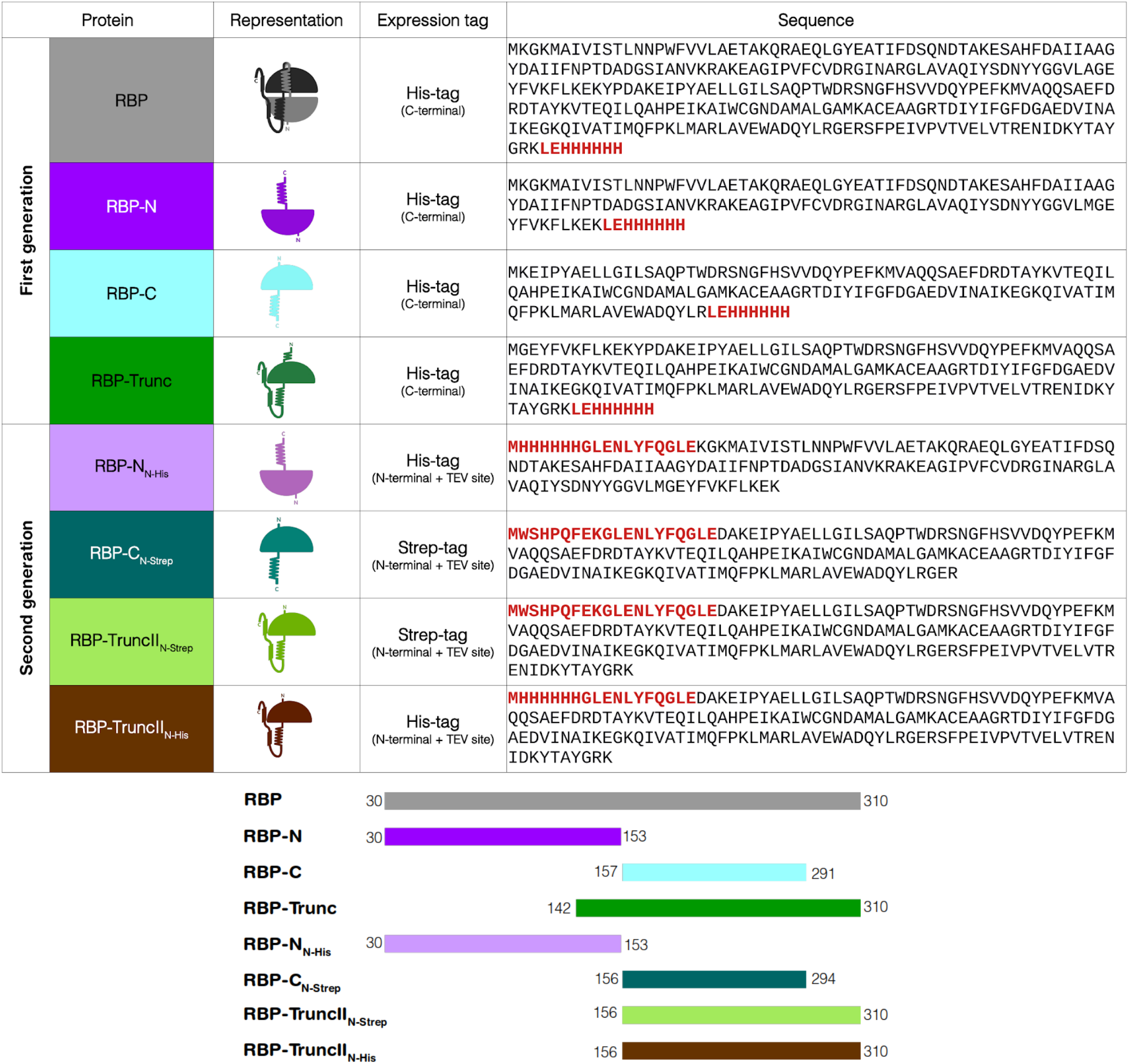
Amino acid sequences of the proteins analyzed in this work. Tags used for expression/purification are highlighted in red. Differences in constructs (numbering consistent with uniprot entry Q9X053) as indicated below. RBP-N & RBP-N_N-His_: correspond to the N-terminal lobe (30-153); RBP-C: corresponding to the flavodoxin-like architecture derived from the RBP C-terminal half, vestigial helix on N- and additional elements on C-terminus removed (157-291); RBP-Trunc: derived from the alternate initiation of translation at M142 (142-310); RBP-TruncII_N-His/N-Strep_: corresponds to truncated construct, with the vestigial helix at the new N-terminus removed (156-310); RBP-C_N-Strep_: corresponds to the C-terminal half, additional residues added of C-terminus (156-294).

**Table S2.**
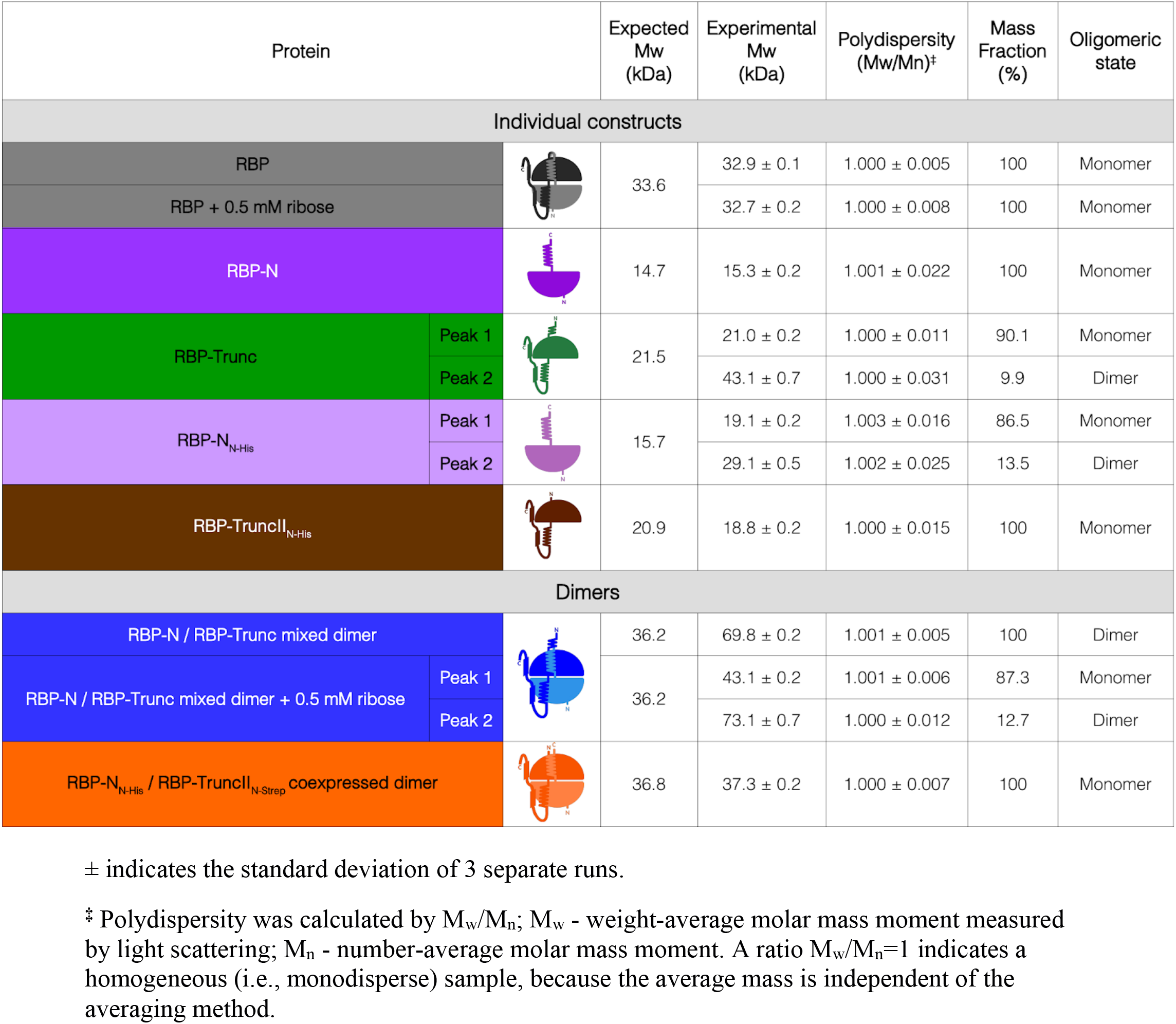
Obtained values from the SEC-MALS measurements for the different RBP constructs.

**Table S3.**
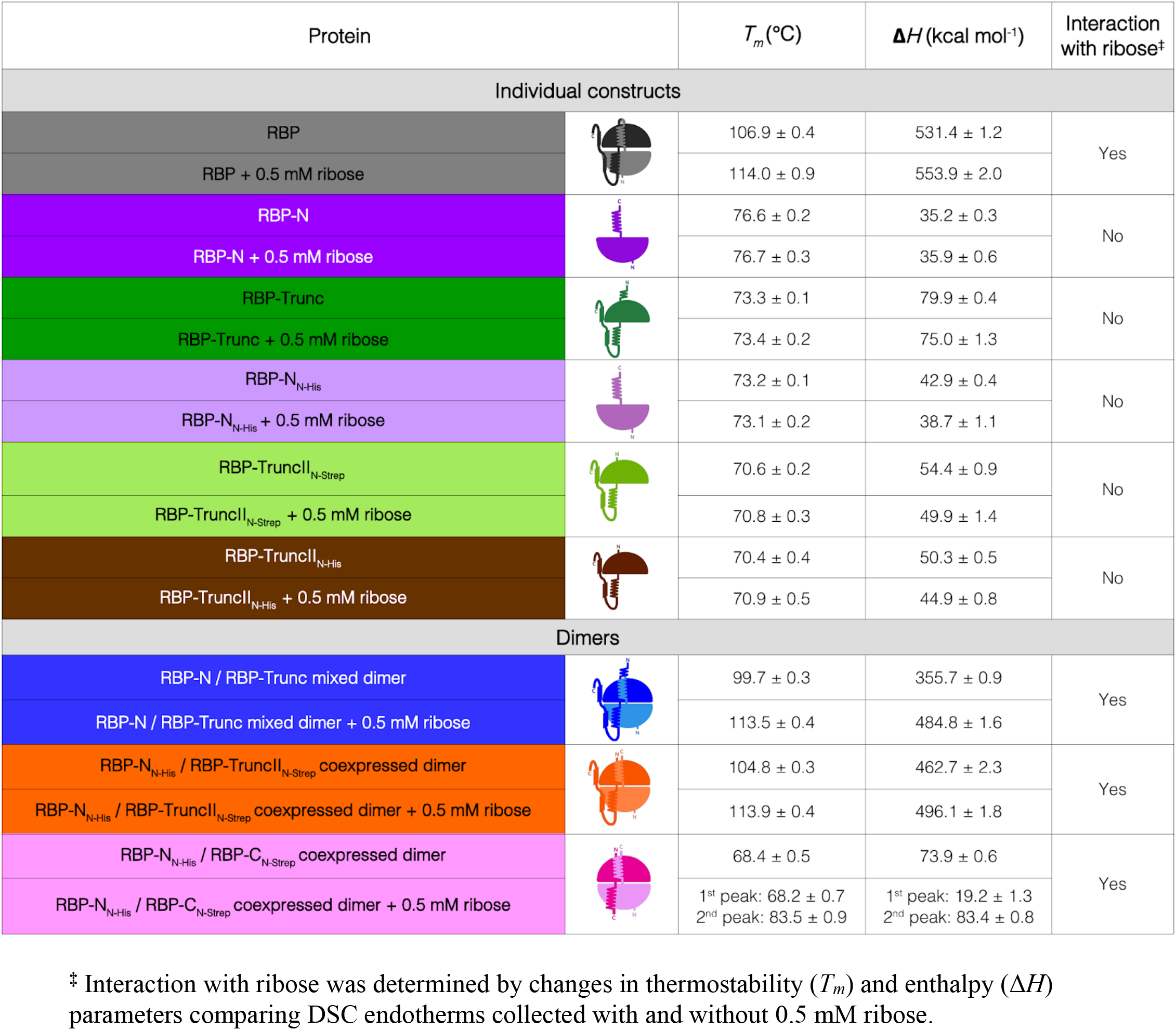
DSC thermodynamic parameters (*T_m_* and Δ*H*) for the different RBP constructs in absence and presence of ribose.

| Protein | | $T_m$ (°C) | $\Delta H$ (kcal mol<sup>-1</sup>) | Interaction with ribose<sup>‡</sup> |
| --- | --- | --- | --- | --- |
| Individual constructs |  |  |  |  |
| RBP |  | 106.9 ± 0.4 | 531.4 ± 1.2 | Yes |
| RBP + 0.5 mM ribose |  | 114.0 ± 0.9 | 553.9 ± 2.0 |  |
| RBP-N |  | 76.6 ± 0.2 | 35.2 ± 0.3 | No |
| RBP-N + 0.5 mM ribose |  | 76.7 ± 0.3 | 35.9 ± 0.6 |  |
| RBP-Trunc |  | 73.3 ± 0.1 | 79.9 ± 0.4 | No |
| RBP-Trunc + 0.5 mM ribose |  | 73.4 ± 0.2 | 75.0 ± 1.3 |  |
| RBP-N<sub>N-His</sub> |  | 73.2 ± 0.1 | 42.9 ± 0.4 | No |
| RBP-N<sub>N-His</sub> + 0.5 mM ribose |  | 73.1 ± 0.2 | 38.7 ± 1.1 |  |
| RBP-TruncII<sub>N-Strep</sub> |  | 70.6 ± 0.2 | 54.4 ± 0.9 | No |
| RBP-TruncII<sub>N-Strep</sub> + 0.5 mM ribose |  | 70.8 ± 0.3 | 49.9 ± 1.4 |  |
| RBP-TruncII<sub>N-His</sub> |  | 70.4 ± 0.4 | 50.3 ± 0.5 | No |
| RBP-TruncII<sub>N-His</sub> + 0.5 mM ribose |  | 70.9 ± 0.5 | 44.9 ± 0.8 |  |
| Dimers |  |  |  |  |
| RBP-N / RBP-Trunc mixed dimer |  | 99.7 ± 0.3 | 355.7 ± 0.9 | Yes |
| RBP-N / RBP-Trunc mixed dimer + 0.5 mM ribose |  | 113.5 ± 0.4 | 484.8 ± 1.6 |  |
| RBP-N<sub>N-His</sub> / RBP-TruncII<sub>N-Strep</sub> coexpressed dimer |  | 104.8 ± 0.3 | 462.7 ± 2.3 | Yes |
| RBP-N<sub>N-His</sub> / RBP-TruncII<sub>N-Strep</sub> coexpressed dimer + 0.5 mM ribose |  | 113.9 ± 0.4 | 496.1 ± 1.8 |  |
| RBP-N<sub>N-His</sub> / RBP-C<sub>N-Strep</sub> coexpressed dimer |  | 68.4 ± 0.5 | 73.9 ± 0.6 | Yes |
| RBP-N<sub>N-His</sub> / RBP-C<sub>N-Strep</sub> coexpressed dimer + 0.5 mM ribose |  | 1<sup>st</sup> peak: 68.2 ± 0.7<br>2<sup>nd</sup> peak: 83.5 ± 0.9 | 1<sup>st</sup> peak: 19.2 ± 1.3<br>2<sup>nd</sup> peak: 83.4 ± 0.8 |  |

<sup>‡</sup> Interaction with ribose was determined by changes in thermostability ( $T_m$ ) and enthalpy ( $\Delta H$ ) parameters comparing DSC endotherms collected with and without 0.5 mM ribose.

**Table S4.** Data collection and refinement statistics for crystal structures. Statistics for the highest resolution shell are shown in brackets.

| Protein | RBP-N / RBP-Trunc dimer |
| --- | --- |
| PDB ID | 7PU4 |
| Wavelength (Å) | 0.9184 |
| Resolution range | 39.01 – 1.69 (1.75 – 1.69) |
| Space group | P 21 21 21 |
| Unit cell [a, b, c (Å) / $\alpha$, $\beta$, $\gamma$ (°)] | 65.2 84.2 103.8 / 90 90 90 |
| Total reflections | 859968 (81810) |
| Unique reflections | 64629 (6298) |
| Multiplicity | 13.3 (12.8) |
| Completeness (%) | 99.76 (98.39) |
| Mean I/sigma(I) | 14.25 (0.96) |
| Wilson B-factor | 30.6 |
| R-merge | 0.129 (1.837) |
| R-meas | 0.127 (1.196) |
| R-pim | 0.035 (1.104) |
| CC1/2 | 0.999 (0.318) |
| CC* | 1.000 (0.695) |
| Matthews coefficient $V_m$ (Å<sup>3</sup> Da<sup>-1</sup>) | 1.97 |
| Solvent content (%) | 37.7 |
| Protein molecules per asymmetric unit | 4 halves |
| Reflections used in refinement | 64515 (6294) |
| Reflections used for R-free | 2095 (205) |
| R-work | 0.206 (0.460) |
| R-free | 0.240 (0.516) |
| CC(work) | 0.964 (0.612) |
| CC(free) | 0.956 (0.581) |
| Number of non-hydrogen atoms | 4757 |
| macromolecules | 4373 |
| ligands | 0 |
| solvent | 384 |
| Protein residues | 574 |
| RMS(bonds) | 0.008 |
| RMS(angles) | 0.900 |
| Ramachandran favored (%) | 97.88 |
| Ramachandran allowed (%) | 1.77 |
| Ramachandran outliers (%) | 0.35 |
| Rotamer outliers (%) | 0.00 |
| Clashscore | 3.75 |
| Average B-factor | 43.1 |
| macromolecules | 43.0 |
| solvent | 44.6 |
| Number of TLS groups | 4 |

## Notes

### Competing Interest Statement

The authors have declared no competing interest.

